# Consensus tetratricopeptide repeat proteins are complex superhelical nanosprings

**DOI:** 10.1101/2021.03.27.437344

**Authors:** Marie Synakewicz, Rohan S. Eapen, Albert Perez-Riba, Daniela Bauer, Andreas Weißl, Gerhard Fischer, Marko Hyvönen, Matthias Rief, Laura S. Itzhaki, Johannes Stigler

## Abstract

Tandem-repeat proteins comprise small secondary structure motifs that stack to form one-dimensional arrays with distinctive mechanical properties that are proposed to direct their cellular functions. Here, we use single-molecule optical tweezers to study the folding of consensus-designed tetratricopeptide repeats (CTPRs) — superhelical arrays of short helix-turn-helix motifs. We find that CTPRs display a spring-like mechanical response in which individual repeats undergo rapid equilibrium fluctuations between folded and unfolded conformations. We rationalise the force response using Ising models and dissect the folding pathway of CTPRs under mechanical load, revealing how the repeat arrays form from the centre towards both termini simultaneously. Strikingly, we also directly observe the protein’s superhelical tertiary structure in the force signal. Using protein engineering, crystallography and single-molecule experiments, we show how the superhelical geometry can be altered by carefully placed amino-acid substitutions and examine how these sequence changes affect intrinsic repeat stability and inter-repeat coupling. Our findings provide the means to dissect and modulate repeat-protein stability and dynamics, which will be essential for researchers to understand the function of natural repeat proteins and to exploit artificial repeats proteins in nanotechnology and biomedical applications.

**Significance statement:** Repetition of biological building blocks is crucial to modulating and diversifying structure and function of biomolecules across all organisms. In tandem-repeat proteins, the linear arrangement of small structural motifs leads to the formation of striking supramolecular shapes. Using a combination of single-molecule biophysical techniques and modelling approaches, we dissect the spring-like nature of a designed repeat protein and demonstrate how its shape and mechanics can be manipulated by design. These novel insights into the biomechanical and biochemical characteristics of this protein class give us a methodological basis from which to understand the biological functions of repeat proteins and to exploit them in nanotechnology and biomedicine.

## INTRODUCTION

Approximately one third of proteins in the human proteome contain repetitive motifs of varying size and structural composition [1, 2]. Within this group, members of the tandem-repeat protein class stand out due to their striking three-dimensional shapes that arise from the stacking of small secondary structure motifs of 20 to 40 amino acids into either quasi-one-dimensional arrays (solenoids) or doughnut-like shapes (toroids). Examples include tetratricopeptide, ankyrin, HEAT and armadillo repeats. Due to their structural simplicity, repeat proteins have been recognised very early to have tremendous potential for applications in nanotechnology, e.g. as synthetic biomaterials, and in biomedicine as antibody alternatives [3, 4]. Previous experiments employed ensemble biochemical techniques to study the folding of repeat proteins and to address the question how alterations in protein sequence translate to changes in protein stability, dynamics and structure [5–7]. However, after two decades of research, it is not yet clear how exactly sequence, shape, stability and dynamics of individual repeats translate into the particular thermodynamic and mechanical properties of the whole array.

Many all-helical repeat proteins look like springs and have indeed been shown to be flexible molecules with spring-like properties, both of which are thought to be crucial to their biological functions [8–18]. Despite this defining feature, our understanding of the mechanics has been limited to date, because the methodology of choice — atomic force microscopy (AFM) — lacks sensitivity in the low-pN regime relevant for these these *α*-helical proteins. To address the limitations of current mechanical measurements of repeat proteins, we employ optical tweezers, which have been used to directly observe conformational transitions close to equilibrium in the low-pN range [19]. Long-term stability and high time-resolution enable us to simultaneously manipulate single repeat proteins, study their spring-like mechanics, and provide detailed information on their dynamics and equilibrium energetics.

Our research focuses on the tetratricopeptide repeat (TPR), which comprises a helix-turn-helix motif and is found in arrays of 3 to 16 repeats in nature [20, 21]. The packing of the TPR motif results in superhelical structures [22], which means that, of all the different repeat-protein types, TPR arrays most closely resemble a physical spring. The functions of TPR proteins are diverse, ranging from scaffolds of multi-protein assemblies and regulators of cell division to molecular chaperones and mediators of bacterial quorum sensing [21, 23–25]. Consensus-designed TPRs (CTPRs) are good candidates for building ‘made to measure’ proteins, because they form stable arrays and are very amenable to manipulation including loop insertions [26–28]. Their chemical stability has been characterised previously [22, 27, 29–36], whereas, in contrast to other repeat proteins [12, 14, 37–41], their mechanical properties remain unexplored. For the above reasons, we chose TPRs to examine how repeat energetics are connected to both the shape and the mechanics of the superhelix.

To achieve this goal, we rationally re-design the geometry of the TPR superhelix by substituting residues at the repeat interface, and then characterise their biophysical properties using Ising models and single-molecule force spectroscopy. Intriguingly, we find that the force response of CTPRs is very different from that of any other protein reported to date. In particular, their rapid dynamics allow them to unfold and refold at equilibrium over a large range of loading rates. We show that by using force we can access the full energy landscape of the CTPR array, which was previously impossible and which allows us to now accurately determine the effect of the chosen mutations. Collectively, our methodology and findings represent an important first step in formulating and redefining future research directions to link repeat protein structure to function.

## RESULTS

### Design and structure determination of a TPR array with altered superhelical geometry

We based our study of the structure and energetics of TPR proteins on previous research of a consensus TPR protein (referred to as CTPRa) that adopts a superhelical structure within the crystal lattice [22]. This superhelical geometry of TPRs is established through stacking of the A-helix (A_*i*+1_ in Fig. 1A) of a given repeat against both A and B helices of the preceding repeat (A_*i*_ and B_*i*_ in Fig. 1A) [26]. The resulting angles between repeat planes then give rise to the pitch and diameter of the superhelix (Fig. 1B) [13]. Interfaces between repeats are largely formed by bulky hydrophobic residues. Therefore, to create a CTPR variant with altered superhelical geometry, we identified four such interface residues based on sequence conservation in different TPR families [26, 42] and substituted them with polar or small hydrophobic side chains (W4L, Y5N, A10V, Y12R).

**FIG. 1.**
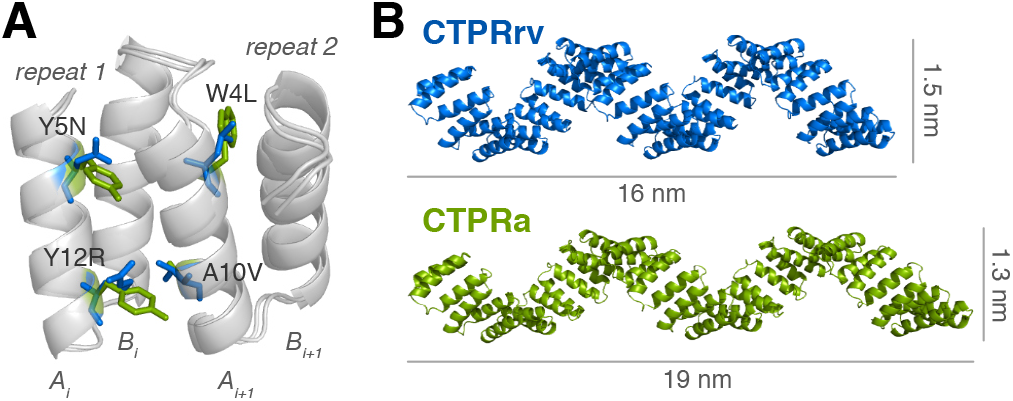
Design of the CTPRrv variant (blue) based on the original CTPRa (green, PDB accession code: 2hyz [22]). (A) Structural representation of two neighbouring CTPRs highlighting the interfacial mutations introduced in CTPRa to form CTPRrv. (B) The slight alteration in repeat packing leads to changes in the diameter and the length of the superhelix, here shown with an array length of 20 repeats.

We determined the structure of a 4-repeat construct of the new variant containing a C-terminal solvating helix, CTPRrv4, by X-ray crystallography to 3.0 Å resolution (see Tab. S3 and Fig. S6). CTPRrv4 crystallised in the P 3_1_ 2 1 space group with two molecules per asymmetric unit. As was observed for CTPRa, the C-terminal solvating helix was not resolved due to a higher preference for end-to-end stacking between molecules in neighbouring asymmetric units [22]. The resolution was sufficient to determine the change in repeat plane angles using the C_*α*_ coordinates: while the twist remained almost unchanged, an increase in curving angle is compensated by a similar decrease in the bending angle (Fig. 1A, Tab. S4). Although these changes may at first appear insignificant in the context of a 4-repeat array, they translate into clear differences in the longer superhelical arrays, leading to a decrease in helix length and increase in helix diameter (Fig. 1B). Furthermore, when compared to other CTPRs, all of which exhibit backbone RMSDs within 0.72 Å, the CTPRrv backbone differs by 1.4 Å relative to the CTPRa backbone.

### Single-molecule force-distance data indicate equilibrium folding of CTPRs

To examine the mechanical folding and unfolding of the CTPR superhelix, we prepared CTPRrv arrays of *N* =3, 5, 10, 20 and 26 and CTPRa arrays of *N* =5 and 9 for force spectroscopy measurements using optical tweezers (Fig. 2A, [43]). Using the Sfp-enzyme, coenzyme-A modifed ssDNA oligos were conjugated to N- and C-terminal ybbR tags on the protein [44]. These protein-DNA chimeras were then attached to micron-sized silica beads using dsDNA handles.

**FIG. 2.**
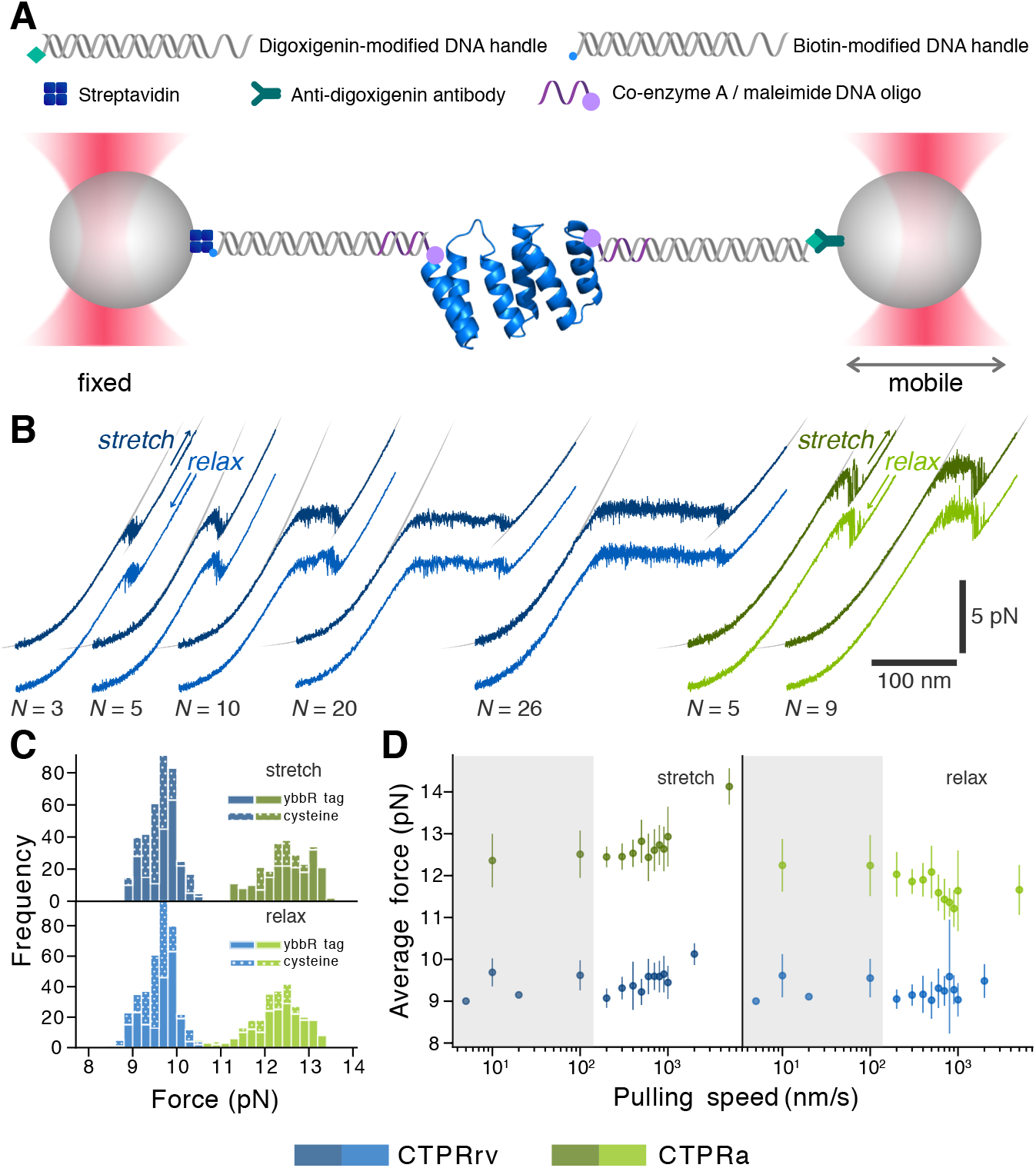
Probing the mechanics of CTPRs using optical tweezers. (A) The protein of interest bears N- and C-terminal modifications that allow the site-specific attachment of ssDNA oligonucletides that can hybridize with single-stranded overhangs of dsDNA handles tethered to micron-sized silica beads. By moving one trap away from the other force is exerted onto the protein-DNA construct resulting in the characteristic stretching of the dsDNA handles, followed by the unfolding of the protein of interest, and finally stretching of the DNA-polypeptide construct. Reversely, if the distance is decreased, the force on the DNA-protein construct relaxes and the protein is allowed to refold. (B) Representative force-distance curves (FDCs) for CTPRrv (blue) and CTPRa proteins (green) of different array length pulled at 10nm/s. The FDCs display a characteristic force ”plateau” and a final ”dip” between the WLC-like stretching of the dsDNA at low forces and the DNA-polypeptide at high forces. The corresponding stretch and relax traces are offset for clarity as they would almost perfectly overlay otherwise. Fits for the DNA WLC and the polypeptide-DNA construct are indicated in grey, and the resulting contour lengths of the unfolded polypeptide are listed in Tab. S2. (C) Histograms of the plateau forces for stretch and relax FDCs of all molecules at 10nm/s and 100nm/s indicate that the transition remains unaffected by the attachment method. (D) The average plateau forces from stretch and relax FDCs at different pulling speeds (shown as mean ± standard deviation to highlight the variation between traces) show only a modest loading rate dependence. This is almost negligible in the rv-type, but a small but clear increase and decrease in unfolding and refolding plateau forces, respectively, can be observed for the a-type. For representative FDCs at different pulling speeds see Fig. S8. The grey shaded area highlights the low pulling speeds at which the unfolding and refolding plateau forces of both repeat types are indistinguishable, and hence the regime in which the respective system is at equilibrium.

For an initial characterization of the CTPR force response, we recorded force-distance curves (FDCs) from stretch-relax cycles at a pulling velocity of 10 nm/s and 100 nm/s (Fig. 2B). The FDCs of all variants display a characteristic plateau region and a subsequent force dip, flanked by the characteristic worm-like-chain (WLC) behaviour of stretching the linker and the linker combined with the unfolded polypeptide. The plateau is preceded by a small and gradual transition that can no longer be described by a WLC. Since the plateau’s length correlates with CTPR array size, we attribute the plateau and dip regions to force-induced unfolding of CTPR repeats. The shape of the unfolding profile is indicative of sequential unfolding of repeats (plateau) until a minimally stable unit is reached, which appears to unravel in a more cooperative manner (dip). Furthermore, we noticed that FDCs from stretch and relax cycles of a single molecule at these loading rates are almost indistinguishable when superimposed. The increased noise levels of plateau and dip indicate very fast unfolding and refolding transitions (Fig. 2B), which, together with the absence of hysteresis, suggests that CTPR unfolding and refolding occurs at equilibrium, i.e. the folding kinetics of the system under investigation are much faster than the pulling speed allowing it to”re-equilibrate instantly”.

The two CTPR variants exhibit the same unfolding and refolding behaviour, albeit at different forces. We estimated the plateau force of each FDC by fitting gaussians to histograms of the respective force data (see SI Methods and Fig. S3). Whereas CTPRa variants unfolded and refolded at a plateau force of ≈12.5pN, the plateau of CRPRrv was significantly lower at ≈9.5pN (Figs. 2B,C), indicating that the introduced mutations have a destabilising effect. Furthermore, to ensure that the intrinsic *α*-helicity of the ybbR-tag [45] did not alter the stability of the repeat arrays, we compared the mechanical and chemical stability of proteins with ybbR-tags to the more commonly used cysteine modifications for maleimide-based attachment strategies (Fig. 2C and Fig. S7). Although a slight stabilisation was observed in chemical denaturation experiments, the ybbR-tag does not discernibly influence the unfolding or refolding plateau forces, the unfolding and refolding energies (beyond the contribution of contour length), or the character of the mechanical response (Fig. 2C).

A systematic screen of pulling velocities ranging from 10 nm/s to 5 μm/s revealed that for both CTPRrv and CTPRa, the average folding and unfolding plateau forces are only marginally affected by the loading rate, leading to a slight increase in hysteresis at pulling speeds of >500 nm/s (Fig. 2D and Fig. S8). This effect is more pronounced in CTPRa arrays than in CTPRrv arrays. It is interesting to note that even at higher pulling speeds not all stretch-relax cycles exhibit hysteresis, leading to a large variation within even a single molecule (e.g. see FDCs collected at 1 μm/s in Fig S9). However, most importantly, we found that there was no significant hysteresis between FDCs from stretch and relax cycles at pulling speeds ≤100nm/s in both repeat types. The absence of a pulling speed-dependent folding/unfolding force is again evidence for rapid equilibrium fluctuations of CTPR subunits in this loading rate regime.

### Averaged force-distance data hint at the folding mechanism of the CTPR arrays

To obtain a more detailed picture of the underlying patters in the equilibrium force response of CTPRs, we binned FDCs of repeated stretch and relax cycles at pulling speeds ≤100nm/s for each individual molecule. Figs. 3A,B show three representative FDCs of each repeat type at the chosen array lengths, overlaid and aligned along force and distance coordinates to avoid the introduction of common instrumental artefacts such as miscalibration of the trap stiffness or the zero distance point. We found the plateau region was not uniformly flat after averaging, as would be expected from other related phenomena such as unconstrained DNA overstretching [46] or the force response of the myosin coiled-coil [47], but rather it showed highly reproducible force oscillations. This pattern was the clearest in the longer arrays of CTPRrv20 and CTPRrv26 which exhibit about 2 and 3 periods, respectively. Therefore, we reasoned that these oscillations arise directly from the structure of the superhelix. In contrast, the characteristic force dip at the end of each FDC was present in all arrays, and we hypothesise that it corresponds to the unfolding of a final stable unit.

**FIG. 3.**
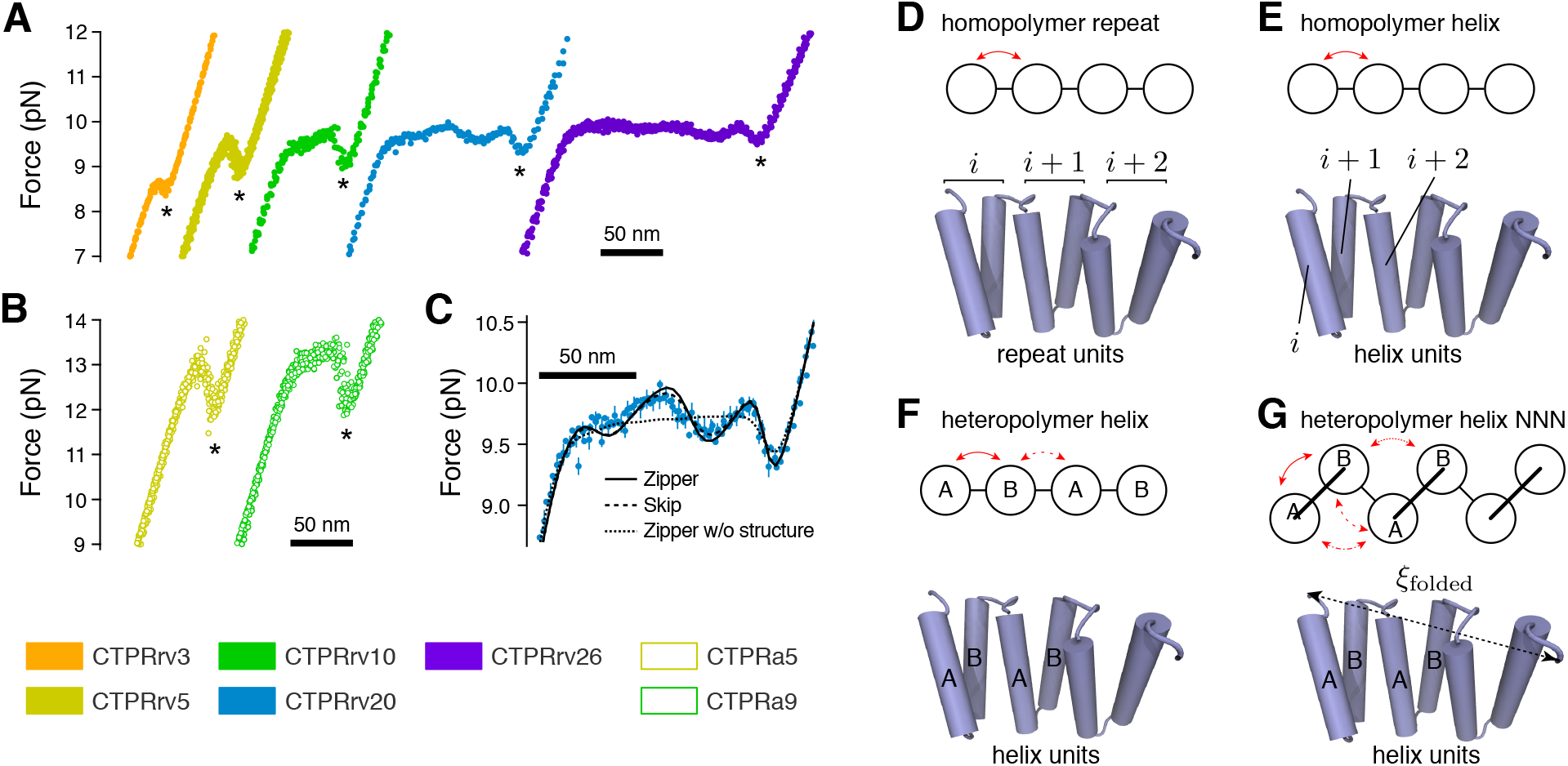
Describing the average force response of CTPRs using Ising models. (A,B) Overlays of aligned equilibrium force-distance curves (FDCs) of CTPRrv and CTPRa variants in which asterisks mark the characteristic force dip after the plateau. (C) Overlay of CTPRrv20 FDCs (cyan) together with fits to the heteropolymer helix Ising models with zipper approximation (solid line), skip approximation (dashed line) and zipper approximation without taking into account structural parameters of the superhelix. In all overlays (A-C), average curves of three representative molecules are shown. (D-G) Different Ising models were tested to describe the folding of TPR proteins. In all models, red arrows indicate the interactions between respective subunits and *ξ*_folded_ represents the end-to-end distance of the folded portion. (D) In the homopolymer repeat model subunits consist of a whole repeats (i.e. two helices). (E) In the homopolymer helix model subunits consist of individual helices that are treated exactly the same. (F) In the heterpolymer helix model the structural repeat is divided into its A and B helices with respective energies. (G) The heteropolymer helix model can be extended to include nearest & next-nearest neighbour interactions (NNN) that may occur e.g. due to structural contacts.

### The force response of CTPRs can be modelled using mechanical Ising analysis

We next set out to model the phenomena observed in our data based on thermodynamic first principles [48–52]. For a given trap distance *d* (see Fig. S4), the total free energy stored in the system, 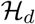 is the sum of the folding free energy of the protein, 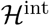, and the mechanical energy for stretching the polymer linkers as well as the traps, 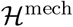:

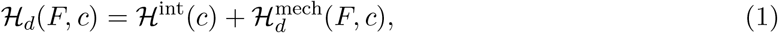

where *F* is the force at that distance and *c* = {*c*_1_,…, *c_N_*} represents a particular configuration of the 2^*N*^-sized conformational space of an *N*-mer. 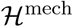 is derived from polymer models of stretching dsDNA and unfolded polypeptide, and the harmonic potential of deflecting the beads from the traps. Theoretical equilibrium FDCs, as are required for fitting data, can subsequently be obtained by numerically calculating the partition function and integrating over all degrees of freedom (see SI Methods for full detail).

Due to the linear stacking of repetitive motifs, we and others have used Ising models to describe the folding energy of repeat proteins [22, 29, 32–36, 53–60]. Hence, we explored various Ising models for 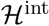. In general, the total energy of an Ising-like system is a linear addition of the energies of the individual units and the interaction energies between nearest neighbours [61, 62]. In repeat proteins, the intrinsic energy of the repeating motif, which can be a single helix or a whole repeat, is considered to be zero when it is unfolded and non-zero when it is folded. The interaction between two neighbouring units (coupling or interfacial stability) then depends on the states they are in: it is only non-zero when two neighbouring repeats are folded but zero when at least one of the neighbours is unfolded.

We first generated models based on a whole repeat (i.e. one A- and B-helix) as the smallest independent protein unit (*homopolymer repeat model*, Fig. 3D). Here, the protein’s internal energy in a given conformation is given by

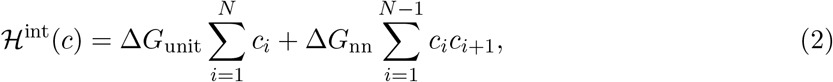

where Δ*G*_unit_ is the folding free energy of a single subunit, Δ*G*_nn_ is the next-neighbour interaction, or coupling energy, between adjacent subunits, and *c_i_* refers to a folded unit. However, this model failed to reproduce the curvature at the transition between the DNA stretch response and the protein unfolding plateau in the CTPRrv10 data (Fig. S5A). Second, we implemented a model that used individual helices as the smallest independent unit based on some of the earliest CTPR folding studies (*homopolymer helix model*, Fig. 3E) [29, 35]. Whereas this model described most of our data much better, we still observed significant deviations for CTPRrv5 molecules (Fig. S5B). Lastly, we expanded the homopolymer helix model to account for the differences between the A- and B-helices within a repeat, and furthermore considered the scenarios in which A and B helices can (i) only interact with their nearest neighbour (*heteropolymer helix model*, Fig. 3F), or (ii) can also form additional, next-nearest neighbour interactions to A- and B-helices in adjacent repeats (*heterpopolymer helix NNN model*, Figs. 1A,3G). In the latter two models, the intrinsic energy is given by

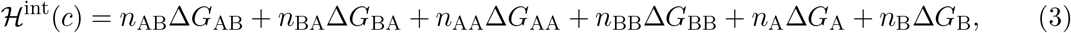

where *n_ij_* count the number of adjacent folded *i* and *j* helices in a configuration *c*, Δ*G_ij_* are the corresponding interaction energies and Δ*G*_A_, Δ*G_B_* are the intrinsic energies of A- and B-helices (see SI Methods for details). The heteropolymer helix model without next-nearest neighbour interactions is recovered when Δ*G*_AA_ = Δ*G*_BB_ = 0. It is important to note that both the heteropolymer helix model and the heteropolymer helix NNN model can be mapped to the repeat model using

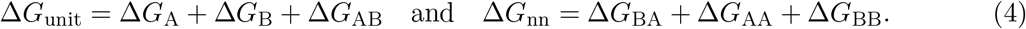

Both heteropolymer helix models were able to describe all data and yielded similar results (Tab. S5). Therefore, we proceeded with the heteropolymer helix model without next-nearest neighbour interaction to avoid over-parametrization. This selection was confirmed by a direct comparison of the models based on the Akaike information criterion (Fig. S5C,D).

All of the above models have exponential 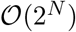 complexity and are computationally too expensive to be applied to CTPRrv20 and CTPRrv26 data. A matrix formalism, which was successfully employed to reduce the complexity of equivalent homopolymer and heteropolymer Ising models in chemical unfolding [56], could not be used to describe the mechanical unfolding because of the non-linear contributions of the linker molecules (DNA and unfolded polypeptide) to the mechanical energy. Instead, we considered alternative simplifications to reduce the conformational space by eliminating extremely unlikely high-energy configurations in two different ways. In the “skip” approximation, all configurations in which one or two folded helices are neighboured by unfolded helices on either side are eliminated. This reduced the complexity from 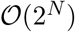 to 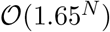 but was still computationally too expensive to fit CTPRrv26 data. However, in the “zipper” approximation, in which unfolding occurs from the end(s), the conformational ensemble is reduced to 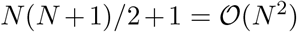 states, and this simplification allowed us to fit data of CTPRrv26 arrays. Notably, the modelled FDCs based on these approximation differed only marginally from the data to which both approximations could be fitted (e.g. see dashed and continuous lines in Figs. 3C, S13A). Furthermore, the resulting energies of the “skip” and “zipper” approximations agreed within error (Tab. 1), and therefore, unless stated otherwise, we used the zipper approximation for all reported values.

**TABLE 1.** Quantitative energetic description of CTPRs. All energies are reported in units of *k_B_T* and *N* refers to the number of repeats. For parameters derived from Ising models, Δ*G*_unit_ is the intrinsic repeat energy, Δ*G*_nn_ the next-neighbour interaction energy, and Δ*G*_tot_ the total energy for an *N*-mer (see Eqs. (4) and (5)). Alternatively, the folding energy can be approximated as the work done by the protein, *W_F_*, from the area under the curve.

| Type | $N$ | Equilibrium denaturation<sup>a</sup> | | | Area<br>$W_F^b$ | Heteropolymer Ising model<br>(Zipper approximation) | | | Heteropolymer Ising model<br>(Skip approximation) | | |
| --- | --- | --- | --- | --- | --- | --- | --- | --- | --- | --- | --- |
| | | $\Delta G_{\text{tot}}$ | $\Delta G_{\text{unit}}$ | $\Delta G_{\text{nn}}$ | | $\Delta G_{\text{tot}}$ | $\Delta G_{\text{unit}}$ | $\Delta G_{\text{nn}}$ | $\Delta G_{\text{tot}}$ | $\Delta G_{\text{unit}}$ | $\Delta G_{\text{nn}}$ |
| rv | 3 | $-13.0 \pm 0.3$ | | | $-22.7 \pm 0.3$ | $-18.4 \pm 0.9$ | $1 \pm 1$ | $-10 \pm 2$ | $-18 \pm 1$ | $1.1 \pm 1.1$ | $11 \pm 2$ |
| | 5 | $-26.1 \pm 0.5$ | | | $-43.4 \pm 0.2$ | $-39.7 \pm 0.4$ | $1.5 \pm 0.3$ | $-11.8 \pm 0.3$ | $-39.7 \pm 0.4$ | $1.5 \pm 0.3$ | $-11.8 \pm 0.3$ |
| | 10 | $-59 \pm 1$ | | | $-91 \pm 1$ | $-87 \pm 3$ | $1.2 \pm 0.3$ | $-11.0 \pm 0.1$ | $-87 \pm 3$ | $1.2 \pm 0.3$ | $-11.0 \pm 0.1$ |
| | 20 | $-125 \pm 2$ | | | $-177 \pm 4$ | $-173 \pm 2$ | $1.0 \pm 0.3$ | $-10.2 \pm 0.2$ | $-173 \pm 2$ | $1.1 \pm 0.2$ | $-10.3 \pm 0.2$ |
| | 26 | $-164 \pm 3$ | | | $-252.7 \pm 0.6$ | $-237 \pm 2$ | $0.5 \pm 0.2$ | $-10.0 \pm 0.3$ | N/A<sup>c</sup> | N/A<sup>c</sup> | N/A<sup>c</sup> |
| | all<sup>d</sup> | | $0.20 \pm 0.05$ | $-6.8 \pm 0.1$ | | | $1.1 \pm 0.2$ | $-11.0 \pm 0.2$ | | $1.3 \pm 0.2$ | $-11.3 \pm 0.3$ |
| a | 5 | $-46 \pm 1$ | | | $-63.7 \pm 0.4$ | $-61.3 \pm 0.6$ | $-2.4 \pm 0.4$ | $-12.4 \pm 0.4$ | $-61.4 \pm 0.6$ | $-2.4 \pm 0.4$ | $-12.4 \pm 0.4$ |
| | 9 | $-91 \pm 2$ | | | $-122.2 \pm 0.9$ | $-118 \pm 2$ | $-1.3 \pm 0.3$ | $-13.3 \pm 0.4$ | $-117 \pm 2$ | $-1.3 \pm 0.3$ | $-13.2 \pm 0.4$ |
| | all<sup>d</sup> | | $-1.0 \pm 0.2$ | $-10.3 \pm 0.1$ | | | $-1.9 \pm 0.3$ | $-12.7 \pm 0.3$ | | $-1.9 \pm 0.3$ | $-12.7 \pm 0.3$ |

<sup>a</sup> Results of the rv-type are based on a global fit to data of arrays with $N = 2, 4, 5, 8$ and $10$ repeats [63], see Fig. S7. Values of the a-type are as reported previously [33]. The errors shown here were propagated from the global fit.
<sup>b</sup> See Eq. (7) in the SI. Values are reported as mean ± s.e.m. for each repeat length.
<sup>c</sup> Values for the heteropolymer model in the skip approximation could not be computed because of limited computational capacity.
<sup>d</sup> Combined value for all repeat lengths.

### Force oscillations are a consequence of the superhelical structure

During the initial rounds of model development we observed that none of our Ising models alone could account for the observed force oscillations in the plateau. At first, the molecular extension of the folded portion was approximated by *ξ*_folded_(*c*) ≈ *n*(*c*)/*N* · *ξ*_max_, where *n*(*c*) is the number of folded subunits in conformation *c* and *ξ*_max_ is the end-to-end distance of the fully folded protein. However, this assumes an arrangement of subunits in the repeat array akin to beads-on-a-string. Although all models based on this assumption for the molecular extension correctly predicted the final dip, they falsely produced a flat plateau (dotted line in Fig. 3C). Only when structural parameters of the superhelix were included to account for the changes in the force vector across the folded remainder of the molecule as it unfolds, was it possible to reproduce these features. Given that our models describe the data well apart from some minor deviations, we can therefore conclude that the structure of CTPR arrays in solution is indeed very similar to that *in crystallo*.

### Comparing intrinsic and interfacial energies between single-molecule and ensemble data

Using our models, we could now determine the intrinsic and interfacial contributions to the free energy, and extrapolated the repeat stabilities and nearest-neighbour coupling from the individual terms of the A and B helices using Equation 4 (Fig. 4A and Tab. 1). Due to the interdependence of the model parameters the data cluster diagonally, i.e. destabilisation of one energetic term is compensated with stabilisation of the other. However, the data of the two variants (filled vs. empty symbols) clearly separate and are independent of attachment method (squares vs. circles). When averaging over the whole data set we found that the interfacial energy 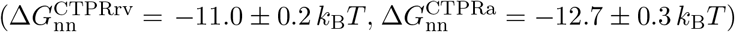 vastly outweighs the intrinsic energy 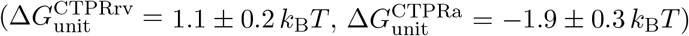 for both repeat types. Next, we determined the total free energy derived from our Ising models using

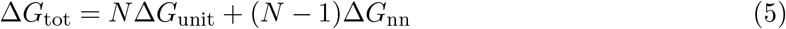

for each variant and array length (diamonds in Fig. 4B and Tab. 1). These values could then be compared to (i) values derived from the area under the curve of FDCs (circles in Fig. 4B and Tab. 1), and (ii) those calculated from ensemble Ising models using Eq. 5 (squares in Fig. 4B, Fig. S7 and Tab. 1). While there is excellent agreement between methods based on single-molecule data, validating our mechanical Ising model approach, energies derived from ensemble measurements severely underestimated the total free energy.

**FIG. 4.**
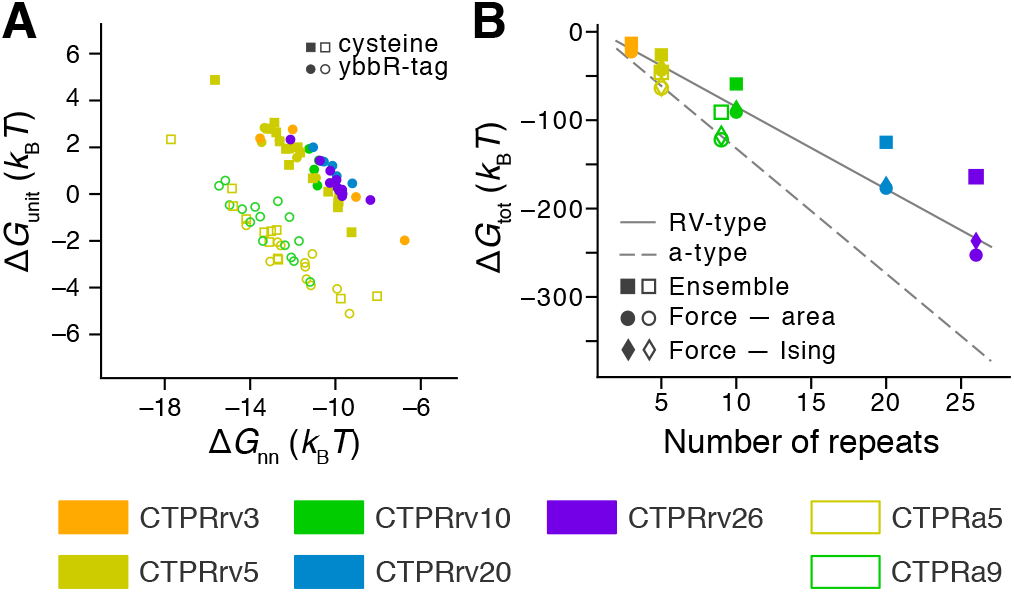
Energetic contributions to the folding free energy derived from single-molecule and ensemble measurements. (A) Intrinsic repeat energy Δ*G*_unit_ and repeat next-neighbour energy Δ*G*_nn_ for individual CTPRrv (filled symbols) and CTPRa molecules (empty symbols) based on the zipper model. For a direct comparison between results derived from skip and zipper approximations see Fig. S9 in the SI. (B) By calculating the total free energy using Eq. 5, a comparison between single-molecule (circles, diamonds) and ensemble measurements (circle) is possible. While the values derived from our Ising models agree very well with those derived from calculating the work done by integration, it is clear that the energies from ensemble experiments deviate significantly. The solid and dashed lines indicate a simple linear regression fits to CTPRrv and CTPRa data obtained from Ising models, respectively, to guide the eye.

### The (un)folding pathway of CTPR arrays

Based on our Ising analyses, we were able to develop a model of the likeliest unfolding (and folding) pathway for CTPR proteins under mechanical load. To this end, we chose to examine results of the skip approximation in more detail to avoid possible bias introduced by the assumption that arrays unfold from the end as it is done in the zipper approximation. For each trap distance we recorded the likeliest conformations ranked by their probabilities. Examples at six different distances along the average unfolding profile of CTPRrv20 are shown in Fig. 5 (and Fig. S10 for CTPRrv26), along with the probabilities of a given configuration and the relative probabilities of individual helices being folded or unfolded. Our data show that unfolding preferentially occurs from the ends in a zipper-like fashion. However, at increasing distances, there are several, almost equally likely conformations that have different segments of folded and unfolded helices. When plotting the probability of being folded for all helices as a function of distance (Fig. S11), we furthermore noticed that unfolding starts with the most C-terminal helix, and then propagates one or two helices at a time from the C-terminus (i.e. A and B helices alternating, or almost in pairs of B_*i*−1_A_*i*_) and one repeat at a time from the N-terminus (i.e. in pairs of A_*i*_B_*i*_).

**FIG. 5.**
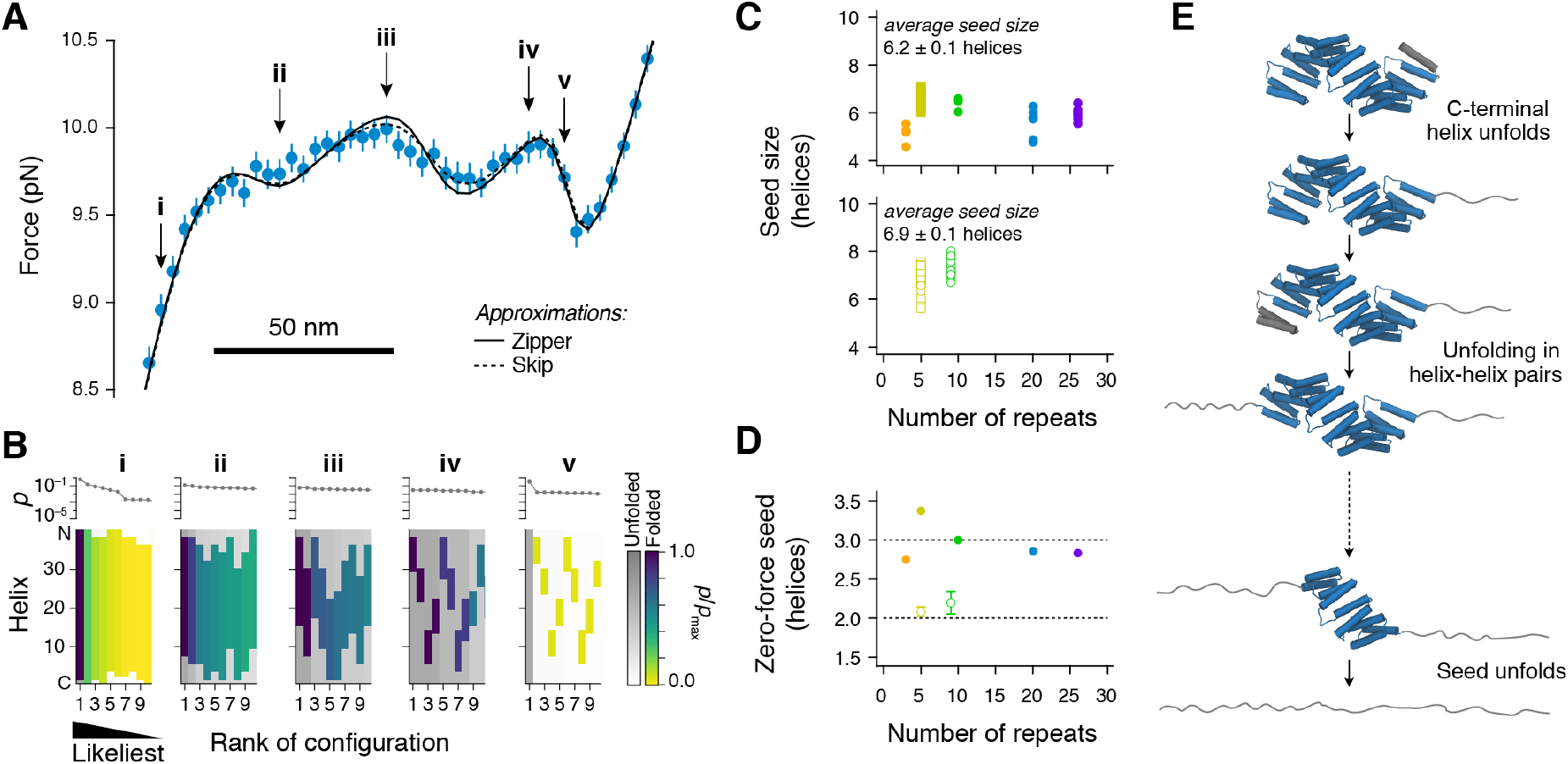
Gaining insights into the folding pathway of CTPR proteins. (A) Average force-distance data for CTPRrv20 fitted to both zipper and skip approximations of the heteropolymer helix model. Roman numerals point to distances for which snapshots of the conformational ensemble are shown in B. (B) From the Ising model, we can extract the ten likeliest configurations at each of the indicated distances in A, ranked according to their probability from highest (index 1) to lowest (shown above the colour maps). Shown are the results for the skip approximation, which does *not* explicitly enforce unfolding from the ends. Coloured bars in the map refer to segments of helices that are folded in a given configuration, with the exact shade giving the relative probability of being folded. Bars in grey-scale on the other hand represent helices that are unfolded, with the exact shade being the corresponding relative probability. Please note, that the N-C-terminal direction is numerically reversed, the C-terminal helix having the index 0 on the y-axis. (C) Average size of the minimally stable folding unit in force experiments for rv- (top) and a-type repeats (bottom). Symbols and colours are the same as those in Fig. 4A. (D) Inferred average minimal stable folding unit in the absence of force for rv-type (filled circles) and a-type variants (empty circles). Error bars show the standard error of the mean. (E) A model for the force-induced unfolding of CTPRs.

Next, we extracted the microscopic information on the characteristic “dip”feature at the end of the plateau region. Traditionally, one would have expected that the dip is caused by the cooperative folding/unfolding of a well-defined minimally stable unit of a certain length that exchanges in a two-state manner with the unfolded state. However, our model indicates that instead of a clear two-state transition, the dip ((iv)–(v) in Fig. 5A,B) represents inter-conversions between a large ensemble of marginally stable conformations of varying size between one another and the unfolded conformation. For example, at position (v) in Fig. 5A, the likeliest conformation is the unfolded state. However, this state exchanges rapidly with an ensemble of conformations with ≈6 to 7 consecutively folded helices that individually are less populated than the unfolded state, but together amount to about 50% of the conformational space (Fig. 5B(v)). We then calculated the average number of folded helices once the fully unfolded conformation reaches a likelihood of 50% to extract the length of the (un)folding “seed”. As expected, the size of this seed was independent of array length, and comprised approximately 6.2 ± 0.1 and 6.9 ± 0.1 helices for the rv- and a-type arrays, respectively (Fig. 5C), which agrees with estimates of the contour length increase extracted from the raw data (Fig. S12). However, it is important to stress that these seeds do not cooperatively exchange with the unfolded conformation in a two-state manner, nor should they be mistaken for the minimal folded unit under zero-force such as one would obtain in ensemble measurements. To get an estimate of the minimal folding unit at zero force, we instead calculated the minimal number of consecutive folded helices needed in order to achieve a negative Δ*G*_tot_ using Eq. (5). The resulting inferred zero-force “seeds” were independent of variant length and were close to 3 helices for CTPRrv and 2 helices for CTPRa (Fig. 5D). Therefore, based on our results, a single CTPRa repeat is weakly stable (−1.9 ± 0.3 *k_B_T*) and may fold transiently on its own. On the contrary, a single CTPRrv repeat is unstable (1.1 ± 0.2 *k_B_T*) and requires the energy from the interface with at least one more helix to fold. These results show the same trend as our ensemble denaturation data, even if they differ numerically.

## DISCUSSION

### CTPR solenoids unzip from both ends under force

CTPRs respond to force in a manner that has never been observed before: they fold and unfold in a single plateau-like transition with regular undulations that ends in a dip. The plateau arises from the asymmetric unzipping of helices or repeats from either end of the repeat array, starting with the C-terminal helix. The very fast equilibrium fluctuations of the C-terminal helix, and possibly also the N-terminal repeat, are beyond the time resolution of our instrument and hence the transition from DNA stretching into the plateau is rather smooth (i.e. “averaged”), particularly in FDCs of the rv-type arrays. The asymmetry of the unfolding pathway arises directly from the structure: (i) the C-terminal B-helix is exposed without any interactions beyond those with the corresponding A-helix of the last repeat of the folded protein/remainder; (ii) the force vector aligns the molecule such that the C-terminal helix can get “un-zipped” while at the N-terminus a whole repeat experiences shear forces; (iii) a B_*i*−1_A_*i*_ repeat is structurally different from an A_*i*_B_*i*_ repeat, leading to different unfolding patterns from either end as the protein is unzipped by force (Fig. S11). This directionality is a natural consequence of the array geometry itself, since repeats at the centre of the array are less likely to unfold than those at the termini, but it is also consistent with hydrogen-deuterium exchange experiments of CTPRs [27, 64] and further studies of other designed and natural repeat proteins. For example, consensus ankyrin repeats were shown to unfold from both ends using chemical denaturation [58] and from one end to the other under force [39]. In contrast, some natural repeat proteins evolved to have repeats (or repeat domains) of significantly different stability, the weakest of which unfold first even if they are located at the centre of the array [65–67].

### To seed or not to seed?

Many globular proteins and also natural repeat proteins have a region or a set of contacts that form a folding nucleus which then “seeds” folding of the remainder. In consensus repeat proteins, such a seed can theoretically form anywhere along the unfolded polypeptide and multiple seeds could form at the same time if the polypeptide is long enough [7]. Consequently, although it is very unlikely that multiple seeds can fold under force, the definition of a seed becomes blurred as we have shown above. However, the average “seed” size of 6 to 7 helices is, intriguingly, consistent with both a folding correlation length of roughly 3 repeats proposed by coarse-grained simulations [7] as well as a folding “nucleus” of 2.5 repeats as concluded from ensemble folding studies of a set of CTPR proteins [32]. In contrast, our estimate of the zero-force seed is much smaller as it only refers to the number of repeating units required for Δ*G* < 0, which, we would like to point out, does not mean that such a structure is fully folded at all times (e.g. a single CTPRa repeat is still 13% unfolded).

### The absence of saw-tooth-like unfolding is a consequence of tether elasticity

To our initial surprise, we did not observe the repeat-by-repeat, saw-tooth unfolding patterns observed for other solenoid repeat proteins, particularly ankyrins. However, using our mechanical Ising model, we increased the effective spring constant of the system connected to the protein (optical trap and linker molecules) to simulate the much stiffer compliance of surface and cantilever in AFM experiments. This modification allowed us to reproduce the characteristic saw-tooth pattern of repeats unfolding one at a time for a consensus ankyrin-repeat protein with five repeats (Fig. S13), highlighting that this behaviour is, at least in part, related to the stiffness of the experimental apparatus rather than an intrinsic characteristic of the protein. These findings raise important questions for future research of repeat proteins under load both *in vitro* and *in vivo* as the context of the set-up or the cellular environment may change how we perceive the force response of the protein of interest.

### The intrinsic and interfacial stabilities are not modulated independently

We set out to create a CTPR array with a different superhelical geometry by making conservative substitutions at the interface between repeats. Our initial assumption was that by changing only interface interactions, we would affect solely the interfacial coupling energy — akin to how the intrinsic repeat stability was modified in a previous study [35]. However, although the chosen mutations did alter the helix packing between repeats more significantly than that within a repeat, ensemble and single-molecule data show that these mutations affected both energetic parameters, i.e. coupling and intrinsic stability. In fact, the mechanical Ising models indicate that the overall lower stability of the rv-type arrays relative to the original CTPR arrays is due to an almost equal destabilization of both intrinsic and interfacial energies (ΔΔ*G*_unit_ ≈ 3 *k_B_T* and ΔΔ*G*_nn_ ≈ 2 *k_B_T*, respectively). We therefore propose that due to the tight interdependence of these two energetic parameters in combination with the structural flexibility of CTPR arrays, most mutations will be compensated by rearrangements in packing geometry and will therefore affect both intrinsic and interfacial energies. Modification of residues on the outside of helices that are not involved in any inter-repeat interactions may simply have a much smaller or no effect on the interfacial coupling, as long as such a modification does not cause any structural rearrangements or changes other side-chain interactions.

### The nature of the unfolded state

Previously, the free energies of unfolding obtained by force spectroscopy were found to be consistent with those measured in ensemble studies or predicted by physical models [50, 51, 68]. Here, however, the differences between energies derived from ensemble and single-molecule force data are much larger than what could be expected from fitting errors and fit parameter correlation. Cortajarena and co-workers have shown that CTPRs can form polyproline-II (PPII) helices at high GdnHCl concentration and consequently are more compact than a random coil [69]. In fact, lattice simulations of simple polymer models were only able to achieve a compaction of the protein similar to that observed experimentally when attractive interactions between PPII helices were taken into account [69]. The presence of such secondary structures in the chemically denatured state can indeed explain the differences found here. In single-molecule force experiments, the formation of PPII helices is prevented and hence the completely unfolded state can be accessed (as judged by the contour length, Tab. S2).

### Force response of the tertiary structure

Previously, several groups used steered MD simulations of natural repeat proteins to show that structural rearrangements at interfaces between repeats allowed the array to stretch as a whole before breaking of the array and unfolding of smaller structural elements occurred at higher forces [10, 11, 16]. However, at this time, we do not have evidence that the CTPR superhelix stretches before unfolding starts at the ends. On the one hand, we did not incorporate an additional term into our models to account for this, primarily to avoid over-parameterization. On the other hand, there is no indication that such a response is of similar compliance to DNA and could therefore be hidden in the DNA response. Indeed, it appears to be the reverse: since DNA parameters compensate for the dimension of the folded construct, we can estimate the end-to-end distances prior to unfolding using a linear regression and obtain values that agree well with our structural data (Fig. S14). Given that we can clearly “see” the superhelix in the force plateau, we believe that the interfaces in CTPR arrays are coupled too strongly to rearrange, a conclusion supported by previous findings that describe packing between CTPRs as rather rigid [70]. This coupling is likely much stronger than the intrinsic stability of the helices and repeats at either end of the array, and therefore unzipping occurs before any stretching can be observed. It remains to be seen how careful modulation of intrinsic repeat stability and interfacial coupling can be used alter the overall stiffness of the tertiary structure to explain the flexibility of several natural repeat proteins observed in both simulations and experiments.

## CONCLUSION

In summary, we have characterised the truly novel force response of a solenoid repeat protein at a quality that is high enough to resolve its tertiary structure. Using Ising models we furthermore have shown how two geometrically distinct CTPRs can differ in their thermodynamic and mechanical properties but still retain the same overall folding profile. Our approach circumvents current drawbacks of ensemble studies, as it only requires data of a single array length and has a clearly defined unfolded state. Hence, we envisage that our findings will be valuable for future endeavours to understand the mechanochemistry of repeat proteins.

## MATERIALS AND METHODS

For a detailed description see the supplementary materials and methods section. In brief, repeat arrays were constructed in the background of a pRSET vector and expressed in *E. coli*. Equilibrium denaturation experiments were performed using guanidine hydrochloride in sodium phosphate buffer pH 6.8, 150mM NaCl in a 96-well plate format [71]. CTPRrv4 with a C-terminal solvating helix was crystallized in a solution containing 0.2 M MgCl_2_, 0.1 M sodium cacodylate pH 6.5 and 50% v/v PEG200. Further details on data collection and processing can be found in the supplementary materials and methods section. Angles between repeat planes were calculated essentially as published previously [13]. Constructs were prepared for force spectroscopy using site-specific modification of either terminal ybbR-tags or cysteine residues [72, 73]. All single-molecule force spectroscopy data was collected on a custom-built instrument [74], processed using custom scripts developed in Igor Pro (WaveMetrics) and further analysed using Igor Pro or Python [75–81]. Theoretical FDCs were calculated using custom C++/CUDA software. Structural representations were generated using PyMol [82] or VMD [83].

## Data availability

All scripts and data are available upon reasonable request. The structure of CTPRrv4 has been deposited in the RCSB PDB with the accession code 7obi.

## Author contributions

A.P.-R. designed the protein variants. A.P.-R., G.F and R.S.E. performed the crystallography and structural refinement. M.S. conducted all other experiments, and M.S. and J.S. analysed the data. D.B. and A.W. provided experimental guidance and maintained the single-molecule equipment. M.S., L.S.I. and J.S. wrote the manuscript. All authors reviewed the manuscript.

## Acknowledgements

MS and AP acknowledge PhD studentship funding from the BBSRC Doctoral Training Partnerships, Cambridge. MS acknowledges additional support from a travel grant provided by the Non-globular Protein Network (COST Action BM1405-39176) and the Eric Reid Fund for Methodology from the British Biochemical Society, and AP from an Oliver Gatty Studentship. RE acknowledges funding of a PhD studentship from AstraZeneca. MR acknowledges funding by the Deutsche Forschungsgemeinschaft (DFG) through SFB863-A2 1111 6624. LSI acknowledges the support of an Senior Fellowship from the Medical Research Foundation of the UK. JS acknowledges the support of the LMU Center for Nanoscience CeNS and a DFG Emmy Noether grant STI673/2-1.

