## Supplementary information for "Consensus tetratricopeptide repeat proteins are complex superhelical nanosprings"

### MATERIALS

All reagents were purchased from Sigma Aldrich, New England Biolabs (NEB), ThermoFisher, Merck or Asco Chemicals unless otherwise stated. 2x yeast tryptone (2xYT) and Lysogeny Broth (LB) Miller were purchased from Formedium. Unmodified DNA oligonucleotides were purchased from Integrated DNA Technologies (IDT) or Sigma Aldrich. Synthetic genes were purchased from IDT. FastDigest restriction enzymes (ThermoFischer), Phusion High-Fidelity DNA polymerase (NEB), and QuickStick Ligase (Bioline, discontinued) or the Anza T4 Ligase Master Mix (Invitrogen) were used for all cloning processes. *E. coli* strains for molecular biology were purchased from Bioline (α-select Competent Cells, Gold/Bronze Efficiency, discontinued) or NEB (NEB 5-alpha Competent *E. coli*, High efficiency). *E. coli* cells for expression were generated in house from C41 cells obtained from the Kommander Lab (MRC-LMB, Cambridge). All constructs were expressed in vectors based on a pRSET backbone (Ampicillin resistance).

### PROTEIN SEQUENCES

The majority of CTPRs used for this study are based on the consensus sequence containing (a) the terminal RS residues arising from the BglII restriction site that is required for constructing longer repeat arrays [1, 2], and (b) the QK mutation for charge balancing of the final repeat protein [3]. The four-repeat construct used for crystallography was purchased as a synthetic gene, and contained the consensus asparagine residues at the repeat termini as well as a solvating helix.

In the following sequences the pre/suffixes c and y identify cysteine and ybbR-tag attachment points for handles.

#### (CTPRrv)$_\mathbf{N}$

MRGSHHHHHHGLVPRGS(AEALNNLGNVYREQGDYQKAIEYYQKALELDPRS)$_N$

#### y(CTPRrv)$_\mathbf{N}$y

MRGSHHHHHHGLVPRGSDSLEFIASKLA(AEALNNLGNVYREQGDYQKAIEYYQKALELDPRS)$_N$DSLEFIASKLA

#### c(CTPRrv)$_\mathbf{N}$c

MRGSHHHHHHNNNNNNNNNNENLYFQGCGS(AEALNNLGNVYREQGDYQKAIEYYQKALELDPRS)$_N$KLC

#### CTPRrv construct used for crystallography

MRGSHHHHHHGLVPRGS(AEALNNLGNVYREQGDYQKAIEYYQKALELDPNN)$_4$AEALNNLGNVQRKQG

**$(CTPRa)_N$**

MRGSHHHHHHGLVPRGS(AEAWYNLGNAYYKQGDYQKAIEYYQKALELDPRS)$_N$

**$y(CTPRa)_Ny$**

MRGSHHHHHHNNNNNNNNNNENLYFQGDSLEFIASKLAGS(AEAWYNLGNAYYKQGD YQKAIEYYQKALELDPRS)$_N$KLDSLEFIASKLA

**$c(CTPRa)_Nc$**

MRGSHHHHHHNNNNNNNNNNENLYFQGCGS(AEAWYNLGNAYYKQGDYQKAIEYYQ KALELDPRS)$_N$KLC

### EXPERIMENTAL METHODS

#### Molecular biology

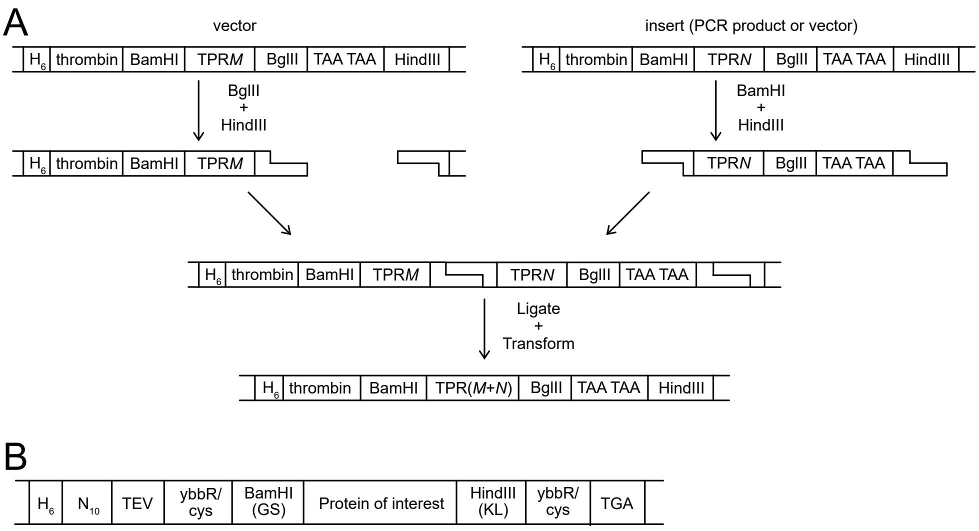


FIG. S1. Schematics illustrating (A) the BamHI-BglII cloning method required to create longer CTPR arrays, and (B) the vector backbone construct developed to facilitate N- and C-terminal modification of proteins for force spectroscopy.

*Mutagenesis*

For Round-the-Horn site-directed mutagenesis (RTH-SDM, [4, 5]), 100 µM primers containing the required mutation/insertion in the overhang were phosphorylated using polynucleotide kinase (ThermoFischer) according to the manufacturer's protocol. Phosphorylated primers were stored at −20 °C until required. The mutation was inserted by PCR, and products were DpnI-digested and gel-purified. About 50 to 100 µg of DNA material was added to 1 µL Anza T4 Ligase Master Mix in a total volume of 4 µL, incubated for 10 to 20 min at room temperature and transformed into *E. coli*. Plasmids were isolated from individual colonies and tested for the presence of the correct mutation/insertion by Sanger sequencing (Eurofins).

*General repeat array construction*

DNA constructs of CTPR proteins in a pRSET backbone were built sequentially from from one, two and four repeat modules using BamHI/BglII cloning as previously described [6]. CTPR repeats are preceded by a BamHI restriction site and followed by a BglII restriction site, double stop codon and HindIII restriction site (Fig. S1). A vector containing $M$ repeats was digested using BglII, HindIII and FastAP Thermosensitive Alkaline Phosphatase (ThermoFisher) according to the manufacturers specifications, and purified using the QIAquick gel extraction protocol. Inserts of up to two repeats were produced by PCR amplification using T7-forward and -terminator sequencing primers. The PCR product was purified according to the QIAquick PCR purification protocol, and digested using BamHI and HindIII followed by heat-inactivation of the enzymes according to the manufacturers specifications. Inserts containing more than two repeats were obtained by restriction digest using BamHI and HindIII and gel extraction. Since BamHI and BglII produce the same 5'-overhangs, the $N$-repeat construct was then ligated directly into the vector using QuickStick (according to the manufacturer's protocol) or Anza T4 ligase (reduced reaction volume as described above), transformed into high efficiency *E. coli* cells, and plasmid purified according to QIAGEN protocols. The whole procedure was repeated until the desired number of repeats was obtained. Using synthetic genes of single repeats, all constructs without tags for DNA attachment were generated this way, and were subsequently used to produce the tagged variants. The construct used for crystallization was obtained as a synthetic gene (Integraed DNA Technologies) and was sub-cloned using the BamHI and HindIII restriction sites. For short arrays (e.g. up to 8 repeats) DNA sequencing could verify the exact number of repeats. Longer arrays were sequenced from both termini to verify the exact cloning boundaries and digested using BamHI and HindIII to determine the number of repeats.

*Construction of yCTPRrv3y and yCTPRrv5y*

Using RTH-SDM, the 11-amino acid ybbR-tags (DSLEFIASKLA) was inserted sequentially between (a) the BamHI restriction site and a TPR, and (b) the BglII site and the stop codons in a construct containing only one repeat (see Fig. S1A, Tab. S1). After digestion with BglII, two and four repeats obtained from BamHI-BglII digests were added at once. The correct orientation of the inserts was identified by restriction digest and Sanger sequencing.

*Construction of yCTPRrv10y, yCTPRrv20y and yCTPRrv26y*

First, ybbR-tags were introduced by RTH mutagenesis directly adjacent to the repeat sequence either N-terminally or C-terminally of a single repeat, giving rise to yCTPRrv1 and CTPRrv1y, respectively. Second, the required number of repeats were added to yCTPRrv1 two or four repeats at a time, resulting in yCTPRrv9, yCTPRrv19 and yCTPRrv25. Last, the C-terminally tagged repeat was added to produce constructs with 10, 20 and 26 that contained both N- and C-terminal ybbR-tags.

*Construction of yCTPRa5y, yCTPRa9y, cCTPRrv5c and cCTPRa5c*

To facilitate ybbR-tagged construct generation, a pRSET vector was modified using RTH-SDM to contain an N-terminal ybbR-tag between TEV cleavage and BamHI restriction sites, and a C-terminal ybbR-tag between the HindIII restriction site and a stop codon (Fig. S1B), Tab. S1). The restriction sites give rise to additional amino acids between the individual ybbR-tags and the protein: GS at the N-terminus and KL at the C-terminus. CTPRa5 and CTPRa10 were assembled in this vector by BamHI/BglII cloning. However, the last two repeats inserted were obtained by a PCR omitting the stop codons (Tab. S1) such that the C-terminal ybbR-tag was in frame. Recombination of CTPRa10 by *E. coli* resulted in a 9-repeat instead of a 10-repeat construct. Since the exact repeat number was irrelevant to our study, we proceeded with this construct.

Proteins containing terminal cysteine residues were created in a similar manner using the same vector but with each ybbR-tag exchanged to a single cysteine (Tab. S1). The CTPRa5 was transferred directly from the corresponding ybbR construct, while the CTPRrv5 had to be re-assembled from a 4-repeat construct fused to a repeat obtained by PCR and without stop codon (Tab S1).

TABLE S1. Sequences of DNA oligonucleotides used for molecular biology.

| **Name** | **DNA sequence (5' → 3')** |
|---|---|
| NybbR Fw | TGCTAGTAAGCTTGCGGCAGAAGCACTGAATAATCTGGG |
| NybbR Rev | ATAAATTCAAGAGAATCGGATCCACGCGGAACCAG |
| CybbR Fw | TGCTAGTAAGCTTGCGTAATAAAAGCTTGATCCGGC |
| CybbR Rev | ATAAATTCAAGAGAATCAGATCTCGGGTCCAGTTCC |
| pRSETa NybbR Fwd | TGCTAGTAAACTTGCGGGATCCGACCTCGAGATCTGC |
| pRSETa NybbR Rev | ATAAATTCAAGAGAATCGCCCTGAAAATACAGGTTTTCGTTG |
| pRSETa CybbR Fwd | TGCTAGTAAACTTGCGTGAGATCCGGCTGCTAACAAAGCCC |
| pRSETa CybbR Rev | ATAAATTCAAGAGAATCAAGCTTCGAATTCCATGGTACC |
| CTPRa2 BamHI Fwd | TGCATGCGGATCCGCCGAGGCGTGGTATAATCTAGG |
| CTPRa2 RS+HindIII Rev | GCATGCATAAGCTTAGATCTTGGGTCGAGTTCTAGGGCC |
| pRSET Ncys Fwd | TGTGGATCCGACCTCGAGATCTGC |
| pRSET Ncys Rev | GCCCTGAAAATACAGGTTTTCGTTG |
| pRSET Ccys Fwd | TGCTGAGATCCGGCTGCTAACAAAGCCC |
| pRSET Ccys Rev | AAGCTTCGAATTCCATGGTACCAGC |
| CTPR_RV1 BamHI Fwd | TGCATGCGGATCCGCAGAAGCACTGAATAATCTGGGTAATGTTTATCG |
| CTPR_RV1 HindIII Rev | GCATGCATAAGCTTAGATCTCGGGTCCAGTTCCAGCGC |

**Protein preparation**

N-terminally $H_6$-tagged CTPR proteins were transformed in C41 *E. coli* and plated on LB Agar containing 100 µg/mL ampicillin. All colonies were used to inoculate 0.5 L of 2xYT media and grown at 37 °C until an optical density between $OD_{600} = 0.6$ and $OD_{600} = 0.8$ was reached, and protein expression was induced with 0.5 mM IPTG over 3-5 hours at 37 °C. After lysis the cell suspension was heated to 70 to 80 °C in a water bath to denature the majority of soluble cellular contaminants. The soluble protein was separated from denatured and insoluble protein fractions by centrifugation for 30 min at 35 000 $\times g$, filtered through a 0.22 µm PES membrane and applied to a 5 mL HisTrap Excel column connected to an Äkta Pure chromatography system and equilibrated in wash buffer (50 mM Tris-HCl pH 7.5, 500 mM NaCl, 20 mM imidazole, SIGMAFAST Protease Inhibitor Cocktail (Sigma), DnaseI (Sigma), Lysozyme (Sigma)). The column was washed with 20 column volumes of wash buffer before proteins were eluted using a high-imidazole buffer (50 mM Tris-HCl pH 7.5, 150 mM NaCl, 300 mM imidazole). All fractions containing protein were pooled, and if necessary, concentrated using a Vivaspin centrifugal concentrator (Sartorius) with the appropriate molecular weight cutoff. The protein was then further purified by size exclusion chromatography using a HiLoad 26/600 Superdex 75 pg or HiLoad 16/600 Superdex 75 pg (GE Healthcare) equilibrated in either Tris or phosphate buffer (50 mM Tris-HCl pH 7.5 or 50 mM sodium phosphate pH 6.8, 150 mM NaCl). Constructs with 10 repeats or more exhibited significant recombination resulting in proteins that had a decreasing number of repeats. Hence, only the first few fractions of the elution peak were pooled for concentration, while >60 % of the fractions had to be discarded.

The CTPRrv4 construct used for crystallography was purified essentially as above but in 50 mM sodium phosphate pH 6.8, 150 mM NaCl based buffers. After elution from the resin with buffer containing imidazole, the protein was dialysed against 50 mM sodium phosphate pH 6.8, 150 mM NaCl for 18 hours, in the presence of thrombin (MP biomedical) to remove the $H_6$-tag from the construct. The protein was further purified using a HiLoad 26/600 Superdex 75 pg column (GE Healthcare) equilibrated in 10 (10 mM HEPES pH 7.5, 150 mM NaCl, and concentrated to 20 mg/mL.

The intact mass of all constructs was confirmed by mass spectrometry.

#### Equilibrium denaturation

Samples of a total volume of 150 µL were prepared in a 96-well format (Greiner, medium-binding), in 50 mM sodium phosphate pH 6.8, 150 mM NaCl with guanidinium hydrochloride (GdHCl) gradients of 0 to 4.5 M (CTPRrv2 and yCTPRrv3y) or 0 to 7 M (all other proteins) [7]. The exact denaturant concentration was calculated using the refractive indices of the native and denaturing buffers. A semi-automatic Hamilton Syringe unit was used to dispense the denaturant gradient. The final protein concentration was adjusted for each construct, depending on repeat type (presence/absence of one tryptophan per repeat) and array length, and ranged from <1 µM (large CTPRrv and all CTPRa constructs) to >11 µM (CTPRrv2). Samples were incubated on an orbital shaker at 25 °C for 2 h. Tryptophan residues were excited at 295 ± 10 nm and fluorescence was monitored at 360 ± 10 nm using a CLARIOStar microplate reader (BMG Labtech). Due to the deletion of tryptophan residues from the CTPRrv variant, tyrosine residues were excited at 280 ± 10 nm and their fluorescence measured at 330 ± 10 nm. The data from 9 reads were averaged and normalised. The resulting fluorescence curve, $F$, was converted to the fraction of folded, $\theta$, or unfolded protein, $1-\theta$, using

$$F = (\alpha_N + \beta_N D)\,\theta + (\alpha_U + \beta_U D)\,(1-\theta) \tag{1}$$

or

$$1-\theta = \frac{-F + \alpha_N + \beta_N D}{\alpha_N - \alpha_U + (\beta_N - \beta_U)\,D}, \tag{2}$$

where $\alpha_N + \beta_N D$ and $\alpha_U + \beta_U D$ describe the base lines at low (native) and high (unfolded) denaturant concentrations. Parameters for the baselines were extracted using a two-state unfolding equation to the whole data set or two separate linear fits to the baselines only.

To extract the intrinsic and interfacial energies ($\Delta G_{\text{unit}}$ and $\Delta G_{\text{nn}}$) a homopolymer repeat Ising model was globally fit to denaturation data of un-tagged constructs with $N$ = 2, 4, 5, 8 and 10 repeats using the PyFolding suite [8], the code of which is based on the formalism developed by Barrick and co-workers [9]. We did not fit a heteropolymer helix model as this would lead to overparametrization (6 free parameters vs. 5 data sets).

### Crystallography

CTPRrv4 at 20 mg/mL was crystallised in JCSG-plus screen, well B10 (0.2 M $MgCl_2$, 0.1 M sodium cacodylate, pH 6.5 and 50% v/v PEG 200, Molecular Dimensions) in sitting drop plates (SwissSci, Molecular Dimensions) with 600 nL droplets in 1:1 and 1:2 ratios of protein to well solution. Crystals were looped and flash frozen without further cryoprotectants. Crystals diffracted to 3.0 Å resolution on beamline I04 at Diamond Light Source (Oxford, UK). The data were processed using autoPROC [10] with the determination of diffraction limits set by a local $I/\sigma I \geq 1.50$. The phase was solved by molecular replacement using a CTPRa4 structure (PDB accession code: 2hyz) with two molecules in the asymmetric unit. Refinements were performed using BUSTER version 2.10.3, [11, 12] and iterative model building in Coot [13]. We conservatively modelled phosphate molecules in the concave face of the TPR superhelix, since this buffer was present during all purification steps prior to size exclusion chromatography. Further details on collection and refinement statistics can be found in Table S3. Models of proteins containing more than 4 repeats were created by symmetry transformation in PyMOL, and missing residues and peptide bonds, e.g. between individual 4-mers, were added using MODELLER [14].

### Calculation of plane angles

Changes in geometry between different repeat protein structures can be measured on two levels: (a) by comparing the whole repeat array (e.g. the superhelical arrangement in the case of TPRs), or (b) by comparing the angular differences between repeat planes. Dimensions of the TPR superhelix were estimated using the "Structure Measurments" tool of UCSF Chimera [15] and 20mer structures. Calculations for obtaining angles between repeat planes were adapted from Forwood *et al.* [16]. In brief, a principal component analysis (PCA) is performed on the $C_\alpha$-atom coordinates of each repeat, omitting the inter-repeat loops, to calculate the principal components (PCs, Fig. S2A) that are orientated along the length (PC1, purple), width (PC2, blue) and depth (PC3, green) of the repeat. As previously reported, curvature is defined as the angle between the respective PC2s of repeats $i$ and $i + 1$ projected onto the plane of repeat $i + 1$, twist is the angle between PC1s projected onto the plane formed by $PC1_{i+1}$ and $PC3_{i+1}$, and lateral bending is the angle of PC3s projected onto the plane formed by $PC1_{i+1}$ and $PC3_{i+1}$ (Fig. S2B). Next, some conventions were introduced to ensure the correct direction (positive or negative) of the angle: (i) PC1 always has the same orientation as the superhelical axis, which is defined by the right-hand-rule from the N- to C-terminal direction of the polypeptide chain [17], (ii) PC3 points into the same direction as a vector from the centroid of repeat $i$ to the centroid of repeat $i + 1$, and (iii) PC2 has the same direction as cross-product of PC3 with PC1. All calculations were performed using custom-written Python scripts with NumPy and Matplotlib extensions [18–21].

### Force spectroscopy experiments

#### *Sample preparation*

Protein-DNA chimeras based on Sfp-mediated conjugation were essentially produced as described previously [22, 23]. Reaction volumes of 50 to 100 µL containing 50 mM HEPES pH 7.5, 10 mM $MgCl_2$, 10 µM ybbR-tagged protein, 20 µM CoA-oligo (Biomers) and 10 µM Sfp-synthase (made in-house, the plasmid was a kind gift from the Gaub Lab at the LMU, Munich) were incubated over-night at room temperature. If necessary, yields pf the desired product were increased by performing the reaction with 40 µM CoA-oligo and 20 µM Sfp-synthase.

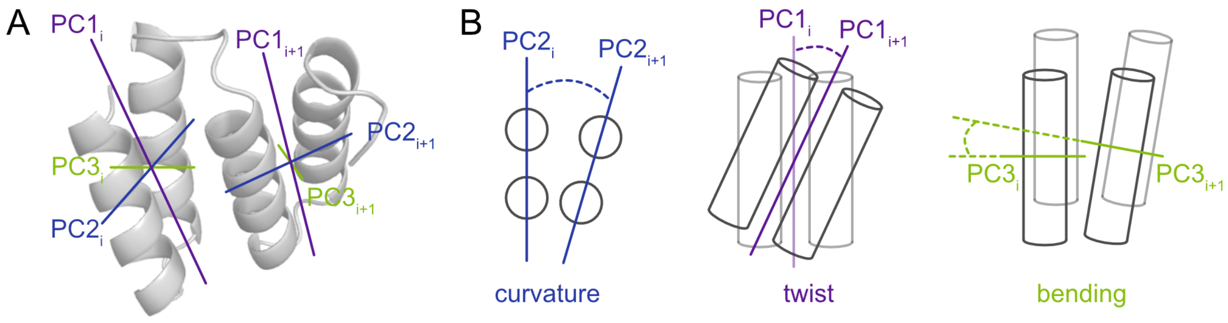


FIG. S2. Visualisation of principal components fitted to repeat planes. (A) Sketch of alignment of PC1-3 with TPR repeats. (B) Schematic representation of how PC1-3 are used to calculate angles for curvature, twist and bending.

Protein-DNA chimeras based on cysteine-maleimide reactions were produced as described previously [24]. In brief, proteins were reduced with a 10-fold excess of TCEP (Sigma Aldrich) for at least 30 min, desalted into phosphate-buffered saline (PBS) using a HiTrap Desalting 5ml (GE Healthcare), and reacted to a 10-fold excess of DBCO-maleimide (Sigma Aldrich) for at least 2 h. After renewed desalting, 10 µM protein was then reacted with 20 µM azide oligo (Integrated DNA Technologies) in 100 µL volumes over-night at 37 °C in an orbital shaker.

Samples were purified using a Superdex 200 10/300 GL (GE Healthcare) or YMC Pack Diol-300 (Yamamura Chemical Research) equilibrated in 50 mM Tris-HCl pH 7.5, 150 mM NaCl. Fractions containing protein conjugated to two oligos were identified by SDS-PAGE, and 4 to 10 µL of those fractions were incubated with 100 to 200 ng biotin- or digoxigenin-functionalised DNA handles at room temperature for at least 30 min. Less than 1 µL of that mixture was added to anti-digoxigenin beads in 10 µL measuring buffer (50 mM Tris-HCl pH 7.5, 150 mM NaCl) and incubated for less than 5 min. Then, 0.5 to 0.7 µL of this mixture were added to 50 µL containing streptavidin beads, an oxygen scavenger system consisting of 0.65% (w/v) glucose (Sigma), 13 U/mL glucose oxidase (Sigma), and 8500 U/mL catalase (Calbiochem). Anti-digoxigenin and streptavidin beads were produced in-house using carboxyl-functionalised 1 µm beads (Bangs Laboratories) [25]. The final mixture was introduced into a home-built chamber that had been blocked with 10 mg/mL BSA for at least 5 min and washed with measuring buffer twice.

*Data acquisition*

All experiments were conducted on a custom-built, dual-beam set up with back-focal plane detection, with both traps having a stiffness of 0.25 to 0.35 pN/nm. An acousto-optical deflector was used to move one bead away from (or towards) the other at speeds ranging between 10 nm/s to 5 µm/s. Bead positions were tracked using a photo-diode detector. Signals were filtered at 50 kHz using an 8-pole Bessel filter, acquired at 100 kHz and downsampled to 20 kHz before storage.

Averaged force-distance curves were obtained from constant-velocity pulling cycles at ≤100 nm/s, where there was no detectable hysteresis by binning, by averaging several stretch FDCs at typically 100 different trap distances.

### DATA ANALYSIS OF RAW FECS AND FDCS

#### Fitting of raw FECs

Force-extension curves (FECs) were fit with

$$F_{\mathrm{eWLC}}(\xi) = \frac{k_{\mathrm{B}}T}{p_{\mathrm{D}}}\left(\frac{1}{4\left(1-\frac{\xi}{L_{\mathrm{D}}}\right)^2} - \frac{1}{4} + \frac{\xi}{L_{\mathrm{D}}} - \frac{F_{\mathrm{eWLC}}}{K}\right) \tag{3}$$

to model the DNA force response [26] and

$$F_{\mathrm{WLC}}(\xi, c) = \frac{k_{\mathrm{B}}T}{p_{\mathrm{p}}}\left(\frac{1}{4\left(1-\frac{\xi}{L_{\mathrm{c}}}\right)^2} - \frac{1}{4} + \frac{\xi}{L_{\mathrm{c}}}\right) \tag{4}$$

to model the unfolded polypeptide [27], where $\xi$ is the extension, $k_{\mathrm{B}}$ is the Boltzmann constant, $T$ the temperature, $p_{\mathrm{D}}$ the persistence length of DNA, $L_{\mathrm{D}}$ the contour-length of the DNA and $K$ its elastic stretch modulus, and $p_{\mathrm{p}}$ and $L_{\mathrm{c}}$ are the persistence and contour length of the protein, respectively. Theoretical and measured protein contour lengths are listed in Tab. S2.

TABLE S2. Expected and measured contour lengths of CTPRa proteins. End-to-end distances $|\Delta\vec{r}|$ are measured between the $C_\alpha$ atoms of the first and last amino acids. The exact length of the ybbR-tags differ between CTPRrv (12 amino acids) and CTPRa (16 amino acids) constructs due to cloning boundaries. All values are in nm. Calculated contour length of the attachment tags are 4.38 nm and 5.84 nm for the ybbR tags of the rv- and a-type proteins, respectively, and 2.19 nm for the cysteine attachments. Measured values are reported as the mean of all molecules and the corresponding standard error.

| **Protein** | **No. molecules** | **Mean no. of traces used for averaging** | $\boldsymbol{L}_{\mathrm{calc}}$ [a] | $\boldsymbol{|\Delta\vec{r}|}$ | $\boldsymbol{L}^*_{\mathrm{calc}}$ [b] | $\boldsymbol{L}_{\mathrm{c}}$ |
|---|---|---|---|---|---|---|
| yCTPRrv3y | 4 | 12 | 41.61 | 3.07 | 34.16 | 30.9 ± 0.4 |
| yCTPRrv5y | 5 | 7 | 66.43 | 4.19 | 57.86 | 56.9 ± 0.7 |
| yCTPRrv10y | 4 | 10 | 128.48 | 7.22 | 117.22 | 116 ± 2 |
| yCTPRrv20y | 7 | 6 | 252.58 | 14.59 | 233.61 | 222 ± 3 |
| yCTPRrv26y | 12 | 5 | 327.04 | 18.86 | 303.8 | 297 ± 1 |
| yCTPRa5y | 11 | 5 | 67.89 | 4.69 | 57.36 | 55.7 ± 0.5 |
| yCTPRa9y | 15 | 6 | 117.53 | 7.65 | 104.04 | 97.8 ± 0.8 |
| cCTPRrv5c | 19 | 7 | 64.24 | 4.19 | 57.86 | 52.2 ± 0.5 |
| cCTPRa5c | 12 | 5 | 64.24 | 4.69 | 57.36 | 52.6 ± 0.6 |

[a] $L_{\mathrm{calc}} = 0.365\,\mathrm{nm} \cdot N_{\mathrm{residues}}$
[b] $L^*_{\mathrm{calc}} = L_{\mathrm{calc}} - |\Delta\vec{r}| - L_{\mathrm{tag}}$

#### Extracting average unfolding and refolding forces

Due to the nature of their unfolding transition, it was not possible to extract the unfolding forces, which traditionally are the force at which a protein or a subdomain unfolds completely, i.e. the force peak. The force data were processed using Igor Pro (Wavemetrics) and analysed further in Python. The data of each force curve were binned into a histogram, giving rise to clear peaks corresponding to the baseline and the unfolding plateau (Figure S3A). The positions of these peaks was extracted from the histogram using a sum of two Gaussian functions and a linear dependence of the background noise on force (force clamping):

$$P(F) = mF + c + a_1 e^{\frac{1}{2}\left(\frac{F-\mu_1}{\sigma_1}\right)^2} + a_2 e^{\frac{1}{2}\left(\frac{F-\mu_2}{\sigma_2}\right)^2}, \quad (5)$$

where $P(F)$ is the probability density of force values, $m$ and $c$ are the slope and intercept of the noise level, and $a$ the scaling factor, $\mu$ the mean and $\sigma$ the standard deviation of the gaussian.

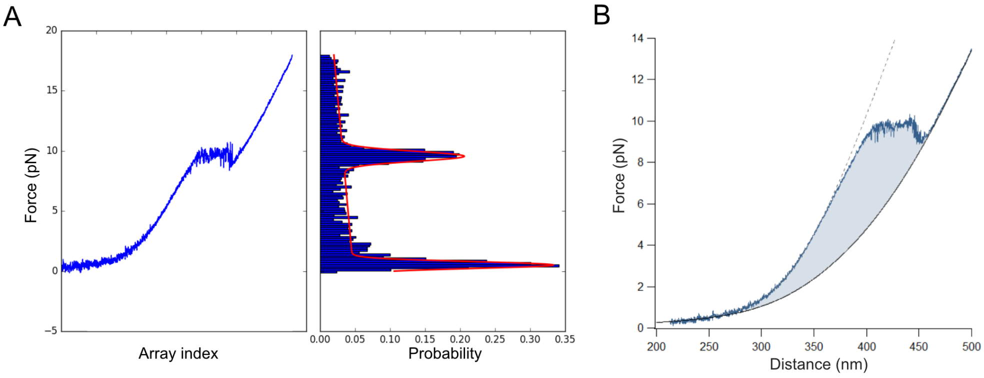


FIG. S3. Calculating the forces and energies of TPR unfolding transitions. (a) The mean unfolding force is extracted by fitting a Gaussian function (red) to a histogram of forces (right) which was derived from the raw data (left, plotted as force against its index array). (b) The non-equilibrium energies of unfolding are simply the area (shaded light blue) between the unfolding curve and the contour of the fully extended construct.

#### Estimating the work done by the trap/protein from constant velocity data

Force-extension curves taken at 10 nm/s and 100 nm/s were fitted with WLC models for both the DNA and fully extended protein. The non-equilibrium energies, or the work done by or on the system, $W$, were then extracted from force-distance curves (FDCs) [28]. The work done on the protein, or the unfolding energy, is simply the difference between the unfolding trace, $U(d)$ and the FDC of the fully extended protein, $C(d)$:

$$W_U = \int_{d_1}^{d_2} U(d)\,\mathrm{d}d - \int_{d_1}^{d_2} C(d)\,\mathrm{d}d, \quad (6)$$

which corresponds to the area between those two curves (Figure S3B). The work done by the protein, or the refolding energy, is the difference between the force response of the unfolded protein and the refolding trace $R(d)$:

$$W_F = \int_{d_1}^{d_2} C(d)\,\mathrm{d}d - \int_{d_1}^{d_2} R(d)\,\mathrm{d}d. \quad (7)$$

### MECHANICAL ISING MODELS

A microscopic conformation $c = \{c_1, \ldots, c_N\}$ of a protein consisting of $N$ subunits was described by a bit-word of length $N$, where ones indicate folded subunits and zeros indicate unfolded subunits. In the case of $N$ subunits there are $2^N$ possible microscopic conformations, e.g. for $N = 3$, $c = \{000, 100, 010, 001, 110, 101, 110, 111\}$.

The full Hamiltonian of the entire system is given by

$$\mathcal{H} = \mathcal{H}_{\text{int}}(c) + \mathcal{H}_{\text{mech}}, \tag{8}$$

where $\mathcal{H}_{\text{int}}(c)$ describes the conformation-dependent internal energy and $\mathcal{H}_{\text{mech}}$ describes the mechanical energy stored in the system.

The energy for mechanically stretching the system consisting of linker and the Hookean spring of the optical trap is

$$\mathcal{H}_{\text{mech}} = \int_0^{d-x} F_{\text{construct}}\,(c,\xi)\,\mathrm{d}\xi + \frac{1}{2}kx^2. \tag{9}$$

In the experimental configuration, the two mechanical parts consisting of dsDNA and unfolded polypeptide are in series (see Fig. S4). Hence, the extension of the full linker consisting of dsDNA and unfolded polypeptide is given by

$$\xi_{\text{construct}}(F,c) = \xi_{\text{eWLC}}(F) + \xi_{\text{WLC}}(F,c) + \xi_{\text{folded}}(c), \tag{10}$$

where $\xi_{\text{eWLC}}$ and $\xi_{\text{WLC}}$ are given by eq. (3) and eq. (11). The extension of the folded protein $\xi_{\text{folded}}$ was assumed to be independent of force, but dependent on the particular configuration $c$ of the protein, i.e. it contained information on the protein structure (see Fig. **??**D). The inverse of eq. (10) yields the force on the construct as a function of length of unfolded polypeptide and total extension $F_{\text{construct}}\,(\xi, c)$.

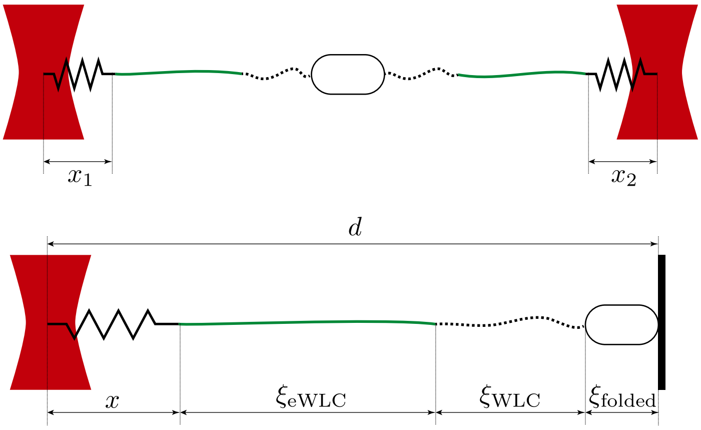


FIG. S4. Lengths and quantities used in the compliance model for a two-bead configuration (top) and the equivalent one-bead configuration (bottom).

The mechanical properties of the dsDNA linker were modelled using Eq. 3, and the mechanical properties of the polypeptide part were modelled using

$$F_{\text{WLC}}\,(\xi, c) = \frac{k_{\text{B}}T}{p_{\text{p}}}\left(\frac{1}{4\left(1-\frac{\xi}{L_{\text{c}}(c)}\right)^2} - \frac{1}{4} + \frac{\xi}{L_{\text{c}}(c)}\right), \tag{11}$$

where $L_{\text{c}}(c) = \left(N - \sum_{i=1}^{N} c_i\right) \cdot L_{\text{aa}} + L_{\text{tag}}$ is the contour length of the unfolded polypeptide when the protein is in conformation $c$, $p_{\text{p}}$ is the persistence length of the unfolded polypeptide, $L_{\text{tag}}$ is the contour length of the attachment tag and $L_{\text{aa}} = 0.365\,\text{nm}$ is the length of a single amino acid.

### Structure information

As highlighted in the main text, the models only accurately described the experimental data when the superhelical nature of CTPR proteins was considered. We incorporated this structural information into eq. 10 by setting $\xi_{\text{folded}}$ to the cumulative end-to-end distance of a folded stretch of helices, as given by the crystal structure.

For example, for a configuration `0111001111`, we set $\xi_{\text{folded}} = \xi_{2\ldots4} + \xi_{7\ldots10}$, where $\xi_{i\ldots j}$ is the crystal-structure end-to-end distance from the start of helix $i$ to the end of helix $j$.

### Interaction models

We considered four different interaction models of subunits and their interactions. For all models, the folded protein extension $\xi_{\text{folded}}(c)$ was obtained from the crystal structure for each possible configuration (see Fig. 3D-G).

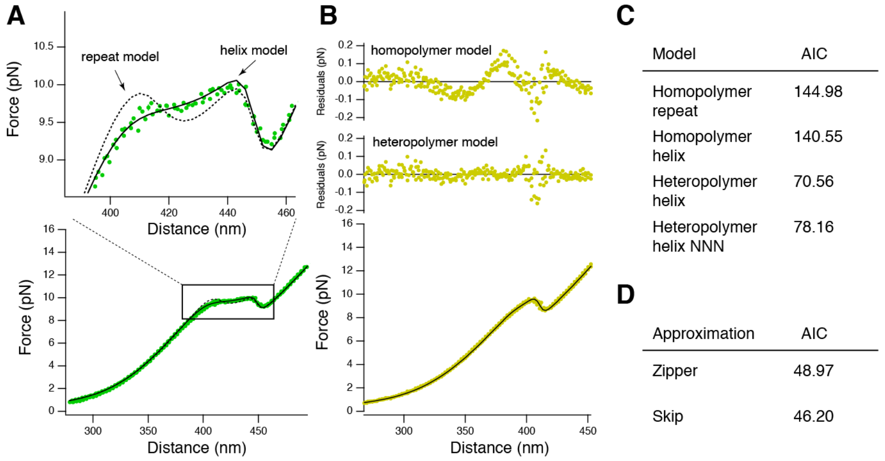


FIG. S5. Model selection. (A) The homopolymer repeat model (dashed line) fails to fit the data for CTPRrv10, while the heteropolymer helix model (continous line) fits well. (B) The homopolymer helix model has higher fit residuals (top) than the heteropolymer helix model (middle) when fitting CTPRrv5 data (bottom). Black line: fit line of heteropolymer helix model. (C) Akaike information criterion (AIC) for the four different interaction models. Reported is the average over all molecules. (D) AIC comparison between the Zipper and Skip approxmations for all molecules ($N = 3, 5, 9, 10, 20$) where the Skip approximation could be fitted.

#### *Homopolymer repeat model*

In the case where repeats are considered subunits, the Hamiltonian for the internal energy of the protein is

$$\mathcal{H}_{\text{int}}(c) = \Delta G_{\text{unit}} \sum_{i=1}^{N} c_i + \Delta G_{\text{nn}} \sum_{i=1}^{N-1} c_i c_{i+1}, \tag{12}$$

where $\Delta G_{\text{unit}}$ is the energy of a folded subunit and $\Delta G_{\text{nn}}$ describes the energy of the next-neighbor interactions between two adjacent folded subunits (Fig. 3D). This is the interaction energy of a one-dimensional Ising model.

*Homopolymer helix model*

The homopolymer helix model is equivalent to the homopolymer repeat model, but subunits consist of helices instead of repeats. Just as for the repeat model, interaction energies only affect next neighbors (Fig. 3E).

*Heteropolymer helix model*

This model takes into account that the two alpha helices in a repeat are different and thus also may be parameterized by different energies. Only next-neighbor energies are allowed. The internal energy is given by

$$\mathcal{H}_{\text{int}}(c) = n_{\text{A}}\Delta G_{\text{A}} + n_{\text{B}}\Delta G_{\text{B}} + n_{\text{AB}}\Delta G_{\text{AB}} + n_{\text{BA}}\Delta G_{\text{BA}}, \tag{13}$$

where $n_{\text{A}}$ is the number of folded A-helices in conformation $c$, $n_{\text{AB}}$ is the number of folded pairs of A and B helices, $n_{\text{BA}}$ is the number of folded pairs of B and A helices, etc (see Fig. 3F).

*Heteropolymer helix nearest & next-nearest (NNN) model*

This model accounts for contacts between adjacent A-A and B-B helices found in the crystal structure and assigns corresponding energies (Fig. 3F). The internal energy of the protein is

$$\mathcal{H}_{\text{int}}(c) = n_{\text{AB}}\Delta G_{\text{AB}} + n_{\text{BA}}\Delta G_{\text{BA}} + n_{\text{AA}}\Delta G_{\text{AA}} + n_{\text{BB}}\Delta G_{\text{BB}} + n_{\text{A}}\Delta G_{\text{A}} + n_{\text{B}}\Delta G_{\text{B}}. \tag{14}$$

Here, $n_{\text{AB}}$ is the number of adjacent folded A and B helices and so on. Unfolded helices are considered to break contacts between next-nearest neighbors, such that a configuration `ABA` would contribute toward $n_{\text{AA}}$, but `A-A` would not.

We note that the both the heteropolymer helix model and the heteropolymer helix NNN model can be mapped to the repeat model when

$$\begin{aligned} \Delta G_{\text{unit}} &= \Delta G_{\text{A}} + \Delta G_{\text{B}} + \Delta G_{\text{AB}} \quad \text{and} \\ \Delta G_{\text{nn}} &= \Delta G_{\text{BA}} + \Delta G_{\text{AA}} + \Delta G_{\text{BB}}. \end{aligned} \tag{15}$$

For all models, the total energy of a protein with $N$ repeats is then

$$\Delta G_{\text{tot}} = N\,\Delta G_{\text{unit}} + (N-1)\,\Delta G_{\text{nn}}. \tag{16}$$

**Calculation of force-distance curves**

Under equilibrium conditions, the mean bead deflection $x$ for a given trap distance $d$ is

$$\langle x\,(d;\eta)\rangle = \frac{\int_x \sum_c x \exp\left(-\frac{\mathcal{H}(c,x;d,\eta)}{k_{\text{B}}T}\right)\mathrm{d}x}{\int_x \sum_c \exp\left(-\frac{\mathcal{H}(c,x;d,\eta)}{k_{\text{B}}T}\right)\mathrm{d}x}, \tag{17}$$

where $\eta$ is the set of model-dependent parameters (e.g. $\Delta G_{\text{nn}}$, $\Delta G_{\text{unit}}$).

Consequently, a force-distance curve (FDC) for a given parameter set $\eta$ can be calculated using

$$F(d;\eta) = \langle x(d;\eta)\rangle \cdot \left(\frac{1}{k_1} + \frac{1}{k_2}\right), \tag{18}$$

where $k_1$ and $k_2$ are the spring constants of the two traps.

#### Calculation of unfolding profile

Similarly, the probability of a subunit $i$ to be folded at a given trap distance $d$ is

$$p_i\left(d;\eta\right) = \frac{\int_x \sum_c \delta_i(c) \exp\left(-\frac{\mathcal{H}(c,x;d,\eta)}{k_\mathrm{B}T}\right) \mathrm{d}x}{\int_x \sum_c \exp\left(-\frac{\mathcal{H}(c,x;d,\eta)}{k_\mathrm{B}T}\right) \mathrm{d}x}, \tag{19}$$

where $\eta$ is the set of model-dependent parameters, and

$$\delta_i(c) = \begin{cases} 1, & \text{if the } i\text{-th bit of word } c \text{ is set} \\ 0, & \text{otherwise} \end{cases}. \tag{20}$$

#### Minimal folding unit under load

To determine the size of the minimal folded unit under force conditions, we first numerically determined $d^* = d \,|\, p(c=0) = \frac{1}{2}$, i.e. the distance at which the unfolded configuration is equally populated as all other configurations, where

$$p\left(c\right) = \frac{\int_x \exp\left(-\frac{\mathcal{H}(c,x;d,\eta)}{k_\mathrm{B}T}\right) \mathrm{d}x}{\int_x \sum_{c'} \exp\left(-\frac{\mathcal{H}(c',x;d,\eta)}{k_\mathrm{B}T}\right) \mathrm{d}x}. \tag{21}$$

The minimal folded unit was then calculated as the mean number of folded subunits of all other configurations $c \neq 0$, weighted by their population.

#### Minimal folding unit in the absence of load

We define the minimal folding unit in the absence of load as the minimal amount of subunits that are necessary such that the total energy of the protein becomes negative.

#### Computation and simplification

FDCs were calculated by numerically evaluating eq. (18) using custom-written CUDA software on a GeForce RTX 2080 graphics card (Nvidia). Even though massive parallelization greatly accelerated the computation time, the calculations were still too expensive for long repeat molecules, such as the 26-repeat protein in the helix models with a conformational space size of $2^{52} \approx 5 \times 10^{15}$. We therefore also considered simplified models where we ignored irrelevant (i.e. high-energy) configurations.

##### *Skip approximation*

For the helix models, we excluded all configurations where an individual helix was folded without adjacent folded neighbors (e.g. `010111`), or where two adjacent helices were folded without stabilizing neighbors (e.g. `110111`). These simplifications were in accordance with previous experimental findings that individual repeats are not stable in solution and resulted in a reduction of the computational complexity from $\mathcal{O}(2^N)$ to $< \mathcal{O}(1.65^N)$.

The simplifications allowed us to calculate FDCs for molecules of all repeat lengths. However, the computational cost for the longest molecules was still very expensive (≈60 h per iteration for one FDC with $\approx 4 \times 10^{10}$ configurations of a 26-mer in the Skip approximation) and prevented us from using these approximations in a fit function.

*Zipper approximation*

Therefore, we also considered a zipper approximation, where unfolding always occurs from the ends and configurations such as `11101111` do not exist. This model was of complexity $\mathcal{O}(N^2)$ and could easily be fitted to all molecules.

*Verification*

In practice, we obtained the energy parameters by fitting the zipper approximation to molecules of all repeat lengths. We then verified that FDCs obtained from the Skip approximation, with the same energy parameters, closely reproduced the prediction of the zipper model (see fig. S9A).

The resulting energies for all molecules for which the computation was feasible were identical within errors when comparing the Skip approximation and the zipper approximation. (see Table 1 in the main text).

**Error estimation and propagation**

To determine the errors of the reported energies $\Delta G_{\text{unit}}$, $\Delta G_{\text{nn}}$ and $\Delta G_{\text{tot}}$ (eqns. (15, 16)), we performed model fits to each individual molecule. The reported errors were then calculated by Gaussian error propagation based on the covariance matrix of the individual values of $\Delta G_{\text{A}}$, $\Delta G_{\text{B}}$, $\Delta G_{\text{AB}}$, $\Delta G_{\text{BA}}$, $\Delta G_{\text{AA}}$ and $\Delta G_{\text{BB}}$ and reported as standard error of the mean (SEM) [29].

### SUPPLEMENTARY TABLES

TABLE S3. Data collection, phasing and BUSTER refinement statistics for the CTPRrv4 structure. Values in parentheses are for the outermost shell.

| Parameters and statistics | PDB ID: 7obi |
|---|---|
| **Data collection** | |
| Space group | $P3_1$ 2 1 |
| Unit cell, a, b, c (Å), | 58.912 58.912 189.517 |
| $\alpha$, $\beta$, $\gamma$ (°) | 90.00, 90.00, 120.00 |
| Resolution range, Å | 51.02 - 3.00 (3.11 - 3.00) |
| Total reflections | 16284 (1550) |
| Unique reflections | 8153 (775) |
| Multiplicity | 2.0 (2.0) |
| Completeness, % | 99.6 (96.9) |
| $I/\sigma I$ | 20.0 (1.3) |
| $R_{merge}$ | 0.017 (0.530) |
| $CC_{1/2}$ | 1.000 (0.858) |
| **Refinement** | |
| $R_{work}/R_{free}$, % | 0.226/0.271 |
| Unique reflections used | 8152 |
| R.m.s deviations: | |
| bond lengths, Å | 0.009 |
| bond angles, ° | 0.96 |
| Ramachandran analysis: | |
| Favoured, % | 98.11 |
| Allowed, % | 2.89 |
| Outliers, % | 0.00 |
| Number of atoms | |
| (average B-factor, Å$^2$): | |
| Protein | 2187 (131.04) |
| Ligands | 20 (177.48) |
| Mean/Wilson B-factor, Å$^2$ | 131.46/114.92 |

TABLE S4. Repeat plane angles calculated for both CTPRa and CTPRrv arrays. Values are presented as mean ± s.e.m. of the three repeat interfaces present in the unit cell of the crystal structure, or of the 19 interfaces present in the structure of a 20 repeat model based on symmetry transformation. Cumulative angles are shown to highlight the differences between the repeat types in small and long arrays. Chain A and B of the CTPRrv crystallographic units produced values within error, hence only values for chain A are shown here.

| **Type** | **Number** | **Curvature** [°] | | **Twist** [°] | | **Bending** [°] | |
|---|---|---|---|---|---|---|---|
| | | $\bar{x}$ | $\sum x$ | $\bar{x}$ | $\sum x$ | $\bar{x}$ | $\sum x$ |
| CTPRa | 4 | $28 \pm 1$ | $83 \pm 4$ | $13.07 \pm 0.03$ | $39 \pm 0.12$ | $22.7 \pm 0.4$ | $68 \pm 1.6$ |
| | 20 | | $497 \pm 20$ | | $256 \pm 0.6$ | | $444 \pm 8$ |
| CTPRrv | 4 | $32 \pm 2$ | 95 | $12 \pm 1$ | 37 | $18 \pm 2$ | 55 |
| | 20 | $31.6 \pm 0.6$ | 601 | $11.4 \pm 0.7$ | 217 | $19.1 \pm 0.7$ | 364 |

TABLE S5. Fitted energy parameters in units of $k_{\mathrm{B}}T$. $N$ is the number of repeats (Zipper approximation). Intrinsic repeat energy $\Delta G_{\mathrm{unit}}$ and repeat next-neighbor interaction energy $\Delta G_{\mathrm{nn}}$ (see eq. (15)). $\Delta G_{\mathrm{tot}} = N\Delta G_{\mathrm{unit}} + (N-1)\Delta G_{\mathrm{nn}}$ is the total energy for a n $N$-mer.

| | | Heteropolymer helix model | | | Heteropolymer helix NNN model | | |
|---|---|---|---|---|---|---|---|
| Type | $N$ | $\Delta G_{\mathrm{tot}}$ | $\Delta G_{\mathrm{unit}}$ | $\Delta G_{\mathrm{nn}}$ | $\Delta G_{\mathrm{tot}}$ | $\Delta G_{\mathrm{unit}}$ | $\Delta G_{\mathrm{nn}}$ |
| rv | 3 | –18.4±0.9 | 0.8±1.1 | –10.3±1.5 | –18.4±1.0 | 0.0±1.3 | –9.2±1.4 |
| | 5 | –39.7±0.4 | 1.5±0.3 | –11.8±0.3 | –39.4±0.5 | 1.3±0.3 | –11.5±0.3 |
| | 10 | –87.0±2.7 | 1.2±0.3 | –11.0±0.1 | –87.1±2.6 | 1.3±0.4 | –11.2±0.3 |
| | 20 | –173.3±2.3 | 1.0±0.3 | –10.2±0.2 | –173.7±2.6 | 1.2±0.3 | –10.4±0.3 |
| | 26 | –236.7±2.4 | 0.5±0.2 | –10.0±0.3 | –238.9±2.2 | 0.5±0.2 | –10.1±0.2 |
| | combined | | 1.1±0.2 | –11.0±0.2 | | 1.0±0.2 | –10.8±0.2 |
| a | 5 | –61.3±0.6 | –2.4±0.4 | –12.4±0.4 | –61.6±0.6 | –2.8±0.3 | –11.9±0.4 |
| | 9 | –117.9±1.6 | –1.3±0.3 | –13.3±0.4 | –119.0±1.5 | –1.6±0.4 | –13.1±0.3 |
| | combined | | –1.9±0.3 | –12.7±0.3 | | –2.3±0.3 | –12.4±0.3 |

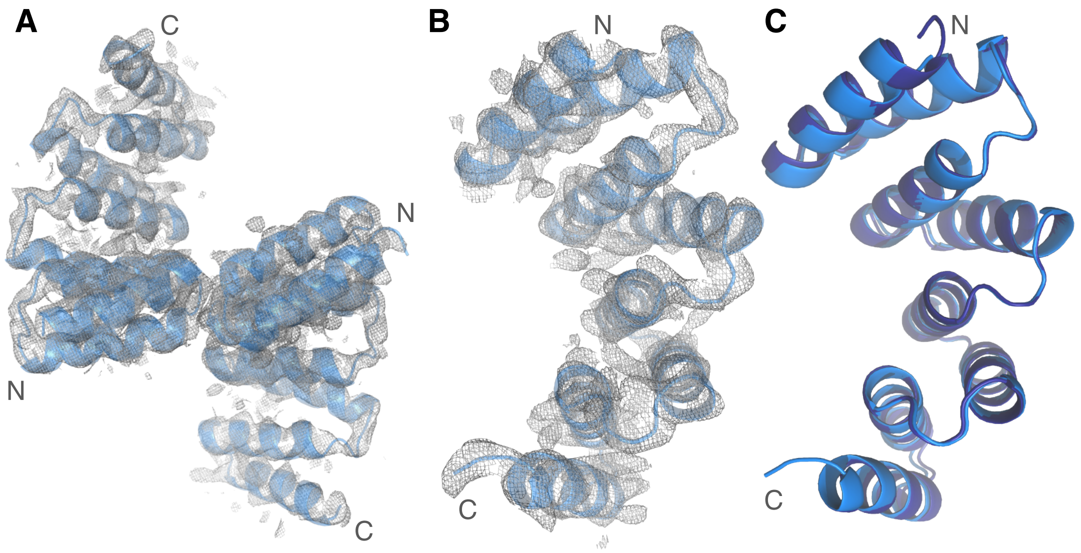


FIG. S6. Crystal structure of CTPRrv. (a) Structures of two macromolecules (marine blue, cartoon representation) present in the asymmetric unit with 2Fo-Fc maps (grey) contoured at 1.5$\sigma$. (b) Zoomed view of chain A, showing clear density for backbone atoms. (c) Structural deviations are minimal between chains A (marine blue) and B (dark blue), an alignment having a backbone RMSD of 0.446 Å.

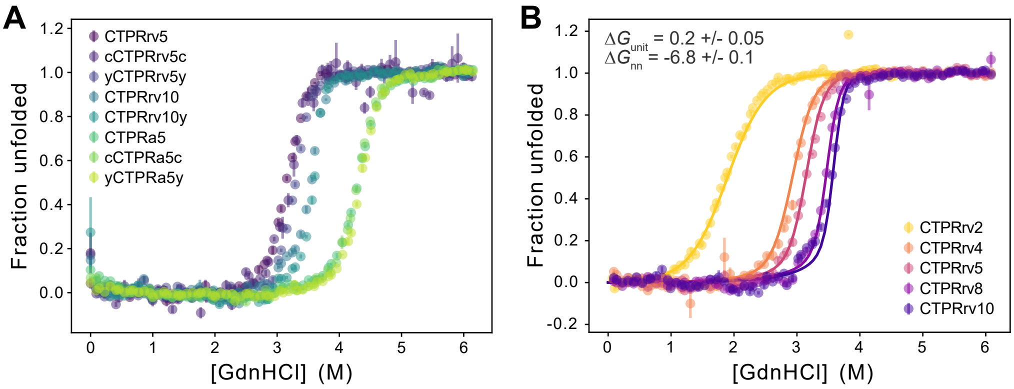


FIG. S7. Equilibrium denaturation data of CTPR arrays using guanidine hydrochloride. (A) Attachment variants of CTPRrv5, CTPRrv10 and CTPRa5 were tested to examine the effect of the added ybbR-tag or cysteine residues at the N- and C-termini. While cysteine modifications did not altered the unfolding profile, the ybbR-tag slightly altered both the transition mid-point and the slope of the transition. We intentionally did not display any fits, since (i) TPRs with more than three repeats clearly deviate from two-state behaviour and (ii) the number of variants was not sufficient to build ensemble heteropolymer Ising models that treated the ybbR-tag as a separate helix with different intrinsic stability and interaction energy at the N- and C-terminal interfaces of the CTPR array. (B) Ensemble Ising models require a global fitting procedure to denaturation data of a series of rv-type arrays with increasing number of repeats. Here, the fits to a homopolymer repeat model with the resulting values for with $\Delta G_{\mathrm{unit}}$ and $\Delta G_{\mathrm{nn}}$ are displayed. A heteropolymer helix model that treated the A- and B-helices different was not fitted as it would result in over-parametrization of the data (6 free parameters versus only 3 used for the homopolymer repeat model). Experiments were performed in technical triplicates in 96-well plate format, and all data are represented as averages with corresponding standard errors.

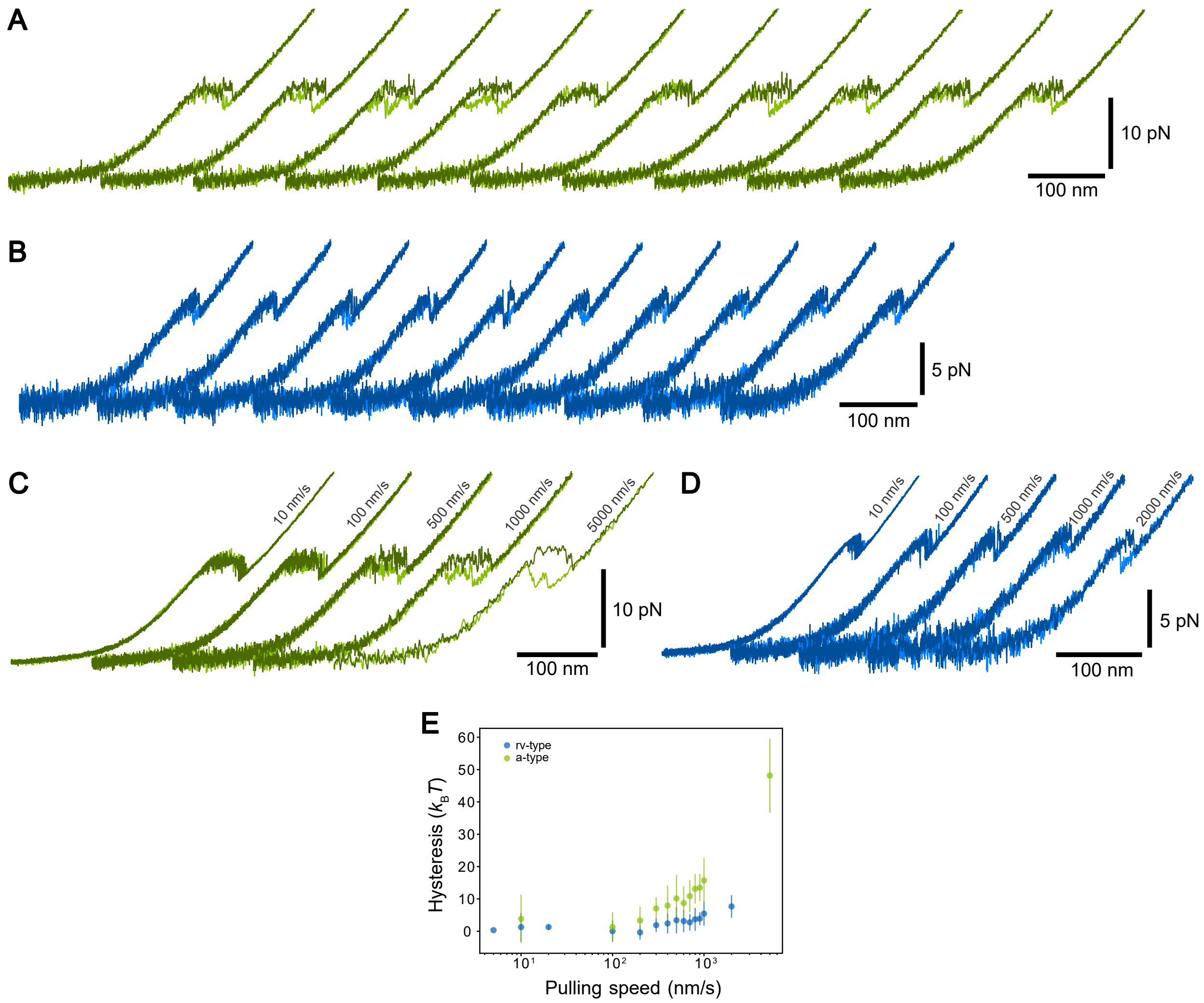


FIG. S8. Hysteresis of CTPR unfolding increases slightly with higher loading rates. (A,B) Consecutive FDCs of one CTPRa9 (green) and one CTPRrv5 molecule (blue) acquired at 1 µm/s highlight the variation observed within a single molecule in the force response at higher pulling speeds. (C,D) Representative traces of the same molecules collected at five different pulling speeds. In all cases the unfolding (darker colours) and refolding traces (lighter colours) are overlaid to highlight the absence or presence of hysteresis. (E) The area under the FDCs was calculated to obtain first estimates of the unfolding and refolding free energies. Using the unfolding and refolding energies it is possible to quantify the hysteresis for individual stretch-relax cycles, here shown as mean with corresponding standard deviations to highlight the increase in variation at the higher pulling speeds. More importantly, this graph clearly shows that hysteresis is negligible for pulling speeds ≤100 nm/s.

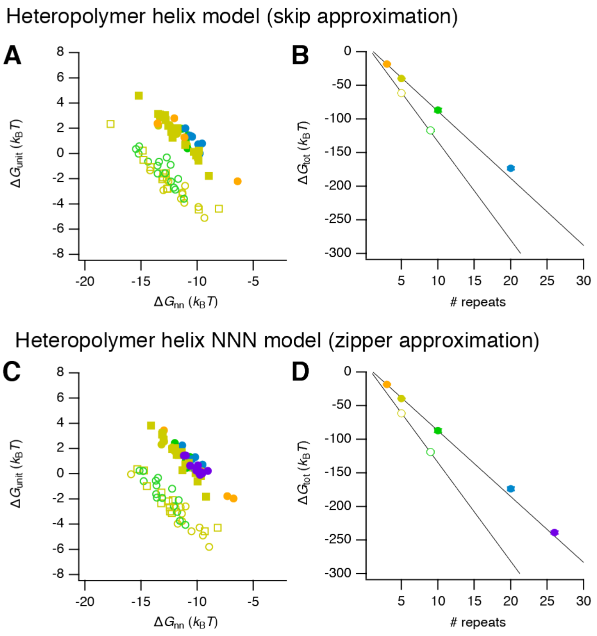

FIG. S9. Zipper and skip approximations of the heteropolymer helix model result in comparable values for $\Delta G_{\mathrm{tot}}$, $\Delta G_{\mathrm{unit}}$ and $\Delta G_{\mathrm{nn}}$. (A,C) Scatter plot of resulting intrinsic repeat energy and next-neighbour interaction energy for each molecule obtained from a heterpolymer helix model with either skip or zipper approximation. Colours/symbols: filled – rv-type, empty – a-type, circles – ybbR attachments, squares - cysteine attachments, colours represent array lengths (see (B,D)). (B,D) The respective total energy $\Delta G_{\mathrm{tot}} = N\Delta G_{\mathrm{unit}} + (N-1)\Delta G_{\mathrm{nn}}$ for each array length of rv-type (filled symbols) and a-type (empty symbols). Error bars represent the SEM, but are too small to be seen.

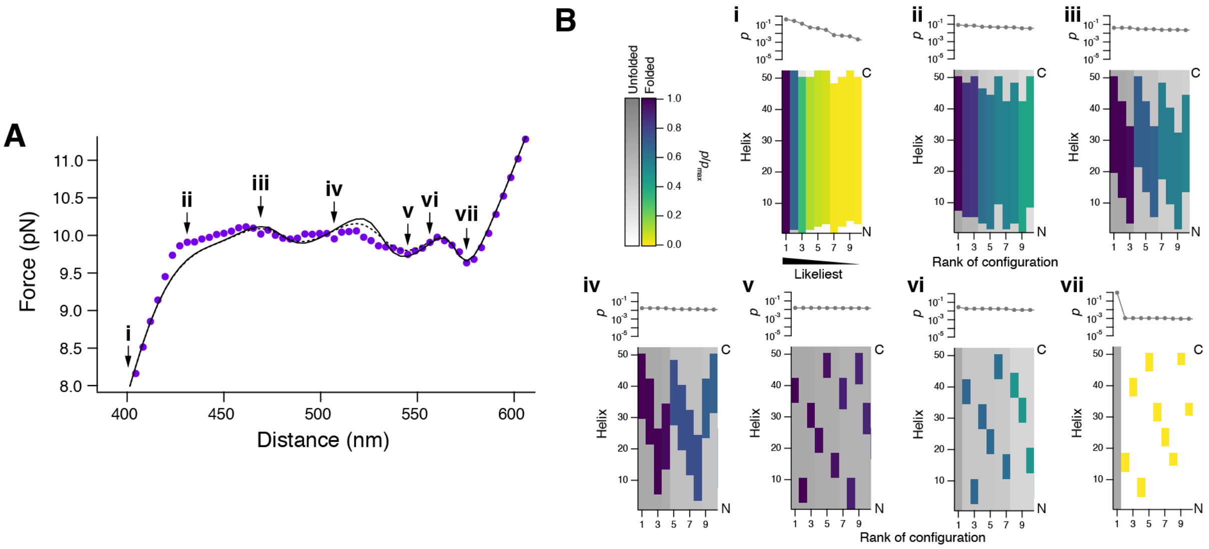

FIG. S10. Predicted unfolding of a 26-repeat protein. (A) Experimental force-distance profile (purple) fitted with a heteropolymer zipper model (continuous black line). Dashed black line: Corresponding prediction from the Skip approximation. Roman letters point to corresponding panels in B. (B) Individual columns represent the ten likeliest configurations at the indicated distances. The likelihood of a particular configuration is shown on top. Color code: Colored stretches are folded, grey/white stretches are unfolded.

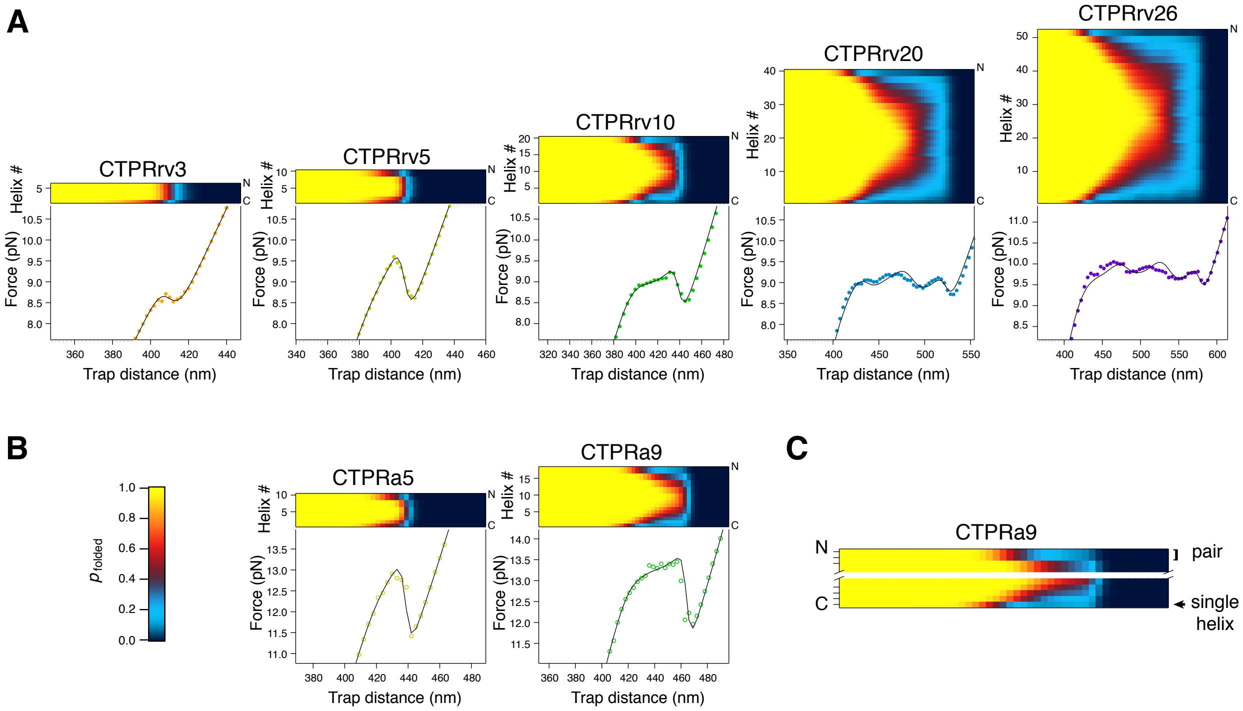


FIG. S11. Unfolding profiles for all measured CTPRrv (A) and CTPRa (B) constructs. Colour maps represent the probability for each helix to be folded as a function of trap distance (please note that indexing proceeds from the C-terminus to the N-terminus in this case). (C) Using a zoom of the CTPRa9 data to exemplify how unfolding starts at the N- and C-termini: in all cases, unfolding starts with the C-terminal helix, and proceeds with the unfolding of (more or less) paired helices from both ends.

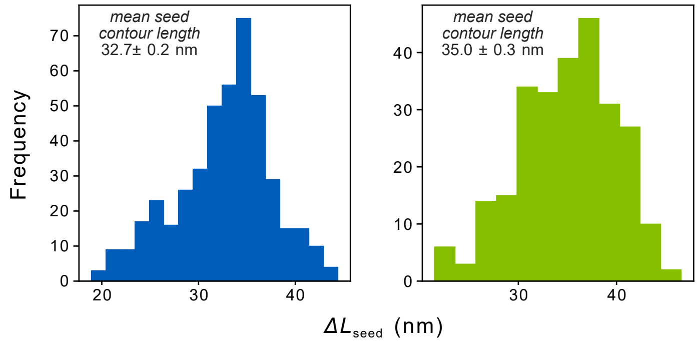


FIG. S12. Contour length histograms of the final "dip" for as measured (roughly) from the end of the plateau to the unfolded contour. Shown are data extracted from FDCs collected at 10 and 100 nm/s of CTPRrv (blue) and CTPR (green). The mean and standard errors for each repeat type are shown. As a reference, the expected contour length increase corresponding to on average 6 helices unfolding is approximately 34 nm, while that of 7 helices unfolding is 38 nm (differences between the two repeat types are less than 1 nm).

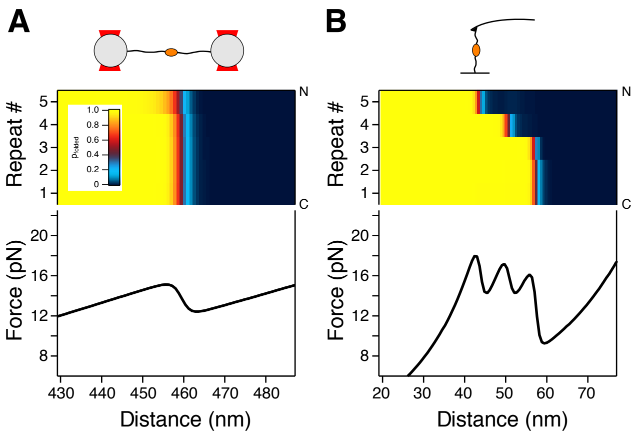


FIG. S13. Simulated FDCs for a consensus ankryin repeat protein in (A) an optical tweezers set-up and (B) under conditions similar to AFM in which linker molecules are much shorter and the protein is tethered between a surface and a much stiffer cantilever. Here we used the structure of the consensus ankyrin $NI_3C$ modelled using the I-Tasser webserver (using all default values [30]), and previously reported values for the energetic parameters of $\Delta G_{\mathrm{unit}} = 5.56\,k_{\mathrm{B}}T$ and $\Delta G_{\mathrm{nn}} = -24\,k_{\mathrm{B}}T$ [31]. Please note, that given this particular structure our results indicate unfolding from the N-terminus to the C-terminus, which is contrary to previous findings.

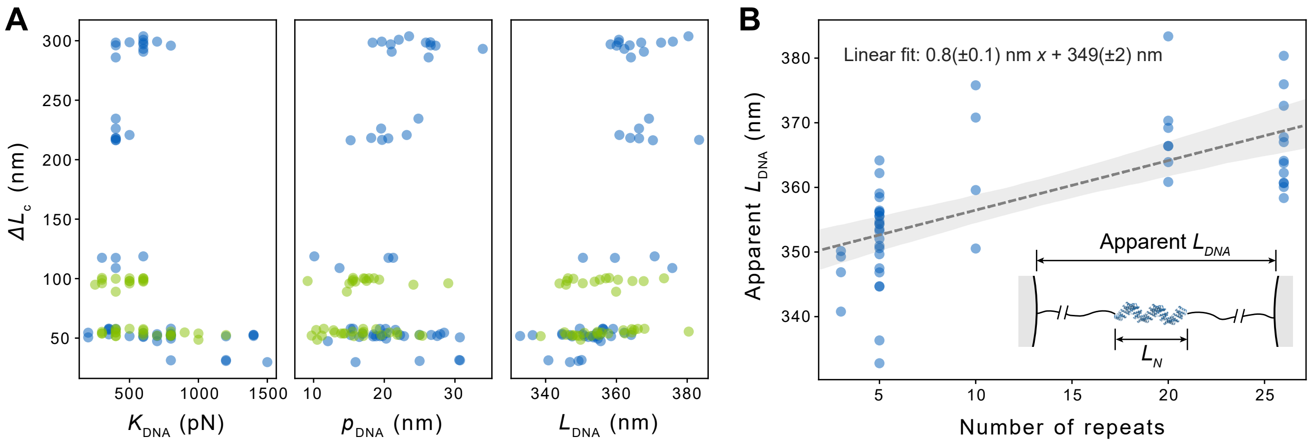


FIG. S14. Fitting DNA-WLCs to raw FDCs, without the explicit use of a protein folding model (Ising or other). (A) There is no indication for a dependence of the protein contour length on any of the DNA parameters. (B) The fitted contour lengths of the tethered constructs are compatible with predictions from the crystal structure. With a rough linear fit, we can estimate an end-to-end distance for CTPRrv20, $L_N \approx 16\,\mathrm{nm}$, based on the increase in the contour length of the full construct (comprising DNA and folded protein) with increasing number of repeats. This value agrees with the crystallographic value (Fig. 1).

[1] E. R. Main, Y. Xiong, M. J. Cocco, L. D'Andrea, and L. Regan, Structure **11**, 497 (2003).

[2] T. Kajander, A. L. Cortajarena, S. Mochrie, and L. Regan, Acta Crystallographica Section D **63**, 800 (2007).

[3] A. L. Cortajarena, T. Kajander, W. Pan, M. J. Cocco, and L. Regan, Protein Engineering, Design and Selection **17**, 399 (2004).

[4] A. Hemsley, N. Arnheim, M. D. Toney, G. Cortopassi, and D. J. Galas, Nucleic Acids Research **17**, 6545 (1989).

[5] S. Moore, "'round the horn site-directed mutagenesis," .

[11] B. G., B. E., B. M., F. C., K. P., P. W., R. P, S. A., S. O.S., V. C., and W. T.O., "Buster," (2020).

[29] I. G. Hughes and T. P. A. Hase, *Measurements and their Uncertainties: A Practical Guide to Modern Error Analysis* (Oxford University Press, 2010).

[30] J. Yang, R. Yan, A. Roy, D. Xu, J. Poisson, and Y. Zhang, Nature Methods **12**, 7 (2015).

[31] S. K. Wetzel, G. Settanni, M. Kenig, H. K. Binz, and A. Plückthun, Journal of Molecular Biology **376**, 241 (2008).
